# JARID2 Inhibition Reprograms Human Hematopoietic Progenitor Cells To Enhance Bone Marrow Transplantation

**DOI:** 10.1101/2025.09.03.672211

**Authors:** Wentao Han, Hassan Bjeije, Hamza Celik, Michael Rettig, Nancy Issa, Andrew L. Young, Yanan Li, Infencia Xavier Raj, Christine R. Zhang, Aishwarya Krishnan, Tyler M. Parsons, Samantha C. Burkart, Jason Arand, Wei Yang, Jeffrey A. Magee, Grant A. Challen

## Abstract

Hematopoietic stem cell transplantation is a common treatment for many blood disorders and can be a life-saving therapy for patients with leukemias, lymphomas and multiple myeloma. Umbilical cord blood (UCB) serves as a valuable source of hematopoietic stem and progenitor cells (HSPCs) for transplantation, particularly for patients lacking a matched donor. However, the limited number of repopulating cells in UCB units restricts its clinical utility. Our prior studies showed that genetic deletion of the polycomb repressive complex 2 (PRC2) co-factor *Jarid2* in mouse multipotent progenitors (MPPs) conveyed ectopic self-renewal capacity. Here, we hypothesized that the function of human HSPCs could be enhanced through *JARID2* inhibition. In this study, we demonstrate that both constitutive and transient knockdown of *JARID2* increases the number and enhances the functionality of human HSPCs both *in vitro* and *in vivo*. This phenotype was distinct from inhibition of *EZH2* in UCB cells, suggesting the mechanism was independent of PRC2 co-factor activity of JARID2. Mechanistically, *JARID2* knockdown promotes a quiescent, long-term self-renewal gene expression program governed by upregulating *STAT1* and characterized by an MHC class II immunophenotype. Analogous to mice, these mechanisms conferred HSC-like potential to human MPPs *in vivo*. Taken together, these findings highlight *JARID2* inhibition as a novel and reversible approach to expand functional UCB-derived HSPCs ex vivo, potentially improving access to stem cell transplantation for a wider patient population.

**One Sentence Summary:** Genetic inhibition of *JARID2* enhances repopulating activity of human hematopoietic stem and progenitor cells *in vivo* via *STAT1* upregulation.

## INTRODUCTION

Bone marrow transplantation (BMT) is a potentially curative therapy for a variety of hematopoietic malignancies including acute and chronic leukemias, Hodgkin and non-Hodgkin lymphoma, myelodysplastic syndromes, and Fanconi anemia (*1*). The functional units of BMT are hematopoietic stem cells (HSCs) contained within the donor graft. The effectiveness of BMT is limited by two primary factors: toxicity (most typically graft-versus-host disease [GvHD]) and donor availability (*2, 3*). Choice of an allogeneic stem cell donor is determined by donor-recipient histocompatibility (based on human leukocyte antigens [HLA]), recipient disease type, recipient condition, and age (*4*). Current sources of stem cell donation from adults are matched siblings (preferred), matched unrelated donors, mismatched unrelated donors, or haploidentical donors. However, only about 30% of patients have a matched sibling available, and the likelihood of finding a matched unrelated donor is low for patients from diverse ethnic backgrounds (*5*). These challenges have led to the rise of umbilical cord blood (UCB) as a donor source. UCB transplantation is now a standard treatment for both children and adults who do not have a matched sibling or unrelated donor. As the hematopoietic cells from newborn babies are immunologically more naïve, there is less risk of GvHD after UCB transplantation and the patient and donor do not need to be as closely matched (*6–8*). UCB transplantation is especially useful for patients of racial and ethnic minorities, and UCB sources are readily available (>700,000 UCB units have been stored for transplantation world-wide)(*9*).

Although UCB transplantation has multiple advantages, it is associated with delayed engraftment and immune reconstitution, thereby leading to greater risks of infection and graft failure compared to adult bone marrow or peripheral blood grafts. This is primarily due to the lower content of HSCs within individual cord blood units. Each UCB unit contains a total HSC dose that is one to two logs lower than the dose contained in adult harvests (*10*). Double-UCB transplantation was implemented to increase the HSC dose and thereby allow adult patients without sufficient-sized single-cord unit to proceed to BMT. Double-UCB transplantation is associated with less graft failure in adult patients, but may produce graft-versus-graft immune complications (*11, 12*). With the increasing use of UCB in BMT, strong efforts are being made to overcome these limitations.

There are intense research efforts aiming to increase UCB-derived HSC numbers or enhance their potency *ex vivo*. The major challenge is to maintain HSC self-renewal potential while promoting the proliferation of cells with repopulation potential. Upon extraction from their native bone marrow (BM) environment, HSCs rapidly lose self-renewal capacity in culture, undergoing substantial changes at both transcriptomic and epigenomic levels. Various strategies have been developed to overcome this limitation in attempts to maintain or expand primary human HSCs *ex vivo*. Notable examples include aryl hydrocarbon receptor antagonists (e.g. StemRegenin-1)(*13*), pyrimidoindole derivatives (e.g., UM171)(*14*), PPAR-γ antagonists(*15*) (e.g., GW9662), and Prostaglandin E2 (PGE2) derivatives (*16–18*), which have all produced mixed success in clinical trials (*19, 20*). Despite these challenges, pursuit of this endeavor remains a worthy goal.

During hematopoietic development, epigenetic modifications are crucial in programming lineage determination and cellular identity (*21*). As epigenetic modifications are reversible and dynamically remodeled during normal development, manipulation of the epigenetic machinery could provide a novel strategy to bias HSC fate decisions and expand populations with reconstituting potential. Recent work in our laboratory has identified a critical role for the Polycomb Repressive Complex 2 (PRC2) co-factor Jarid2 in regulating hematopoietic repopulating potential. Genetic deletion of *Jarid2* in mice confers long-term self-renewal to multipotent progenitor cells (MPPs), the cell population immediately downstream of HSCs in the hematopoietic hierarchy, which possesses only transient repopulating capacity under normal circumstances. As a mechanism, it was found Jarid2 recruits PRC2 to epigenetically silence transcriptional programs associated with HSC cell identity in MPPs through deposition of the H3K27me3 repressive chromatin mark. In the absence of Jarid2 these genes were not epigenetically suppressed in MPPs and sustained expression of genes associated with HSC identity conferred ectopic self-renewal (*22*). A prior study suggested genetic inhibition of *JARID2* may also enhance human HSC function (*23*). With these data in mind, we hypothesized that inhibition of *JARID2* could be utilized to epigenetically reprogram human UCB-derived MPPs with HSC-like potential and enhance BMT.

## RESULTS

### Long-Term Deletion of JARID2 Produces Durable Hematopoiesis Without Transformation

JARID2 is a tumor suppressor in chronic myeloid neoplasms that is lost by hemizygous deletions of the short arm of chromosome 6 during transformation of myeloproliferative neoplasm (MPN) and myelodysplastic syndromes (MDS) patients to secondary acute myeloid leukemia (sAML)(*24–26*). Thus, any potential perturbation of *JARID2* in hematopoietic stem and progenitor cells (HSPCs) could carry inherent risk of neoplastic transformation. To assess the safety of long-term *JARID2* deletion in human HSPCs, we performed CRISPR/Cas9 targeting of *JARID2* in UCB CD34+ cells. 2.5×10^5^ edited cells were transplanted into NSG mice and maintained for one year. Targeting efficiencies were similar for two independent guide RNAs (gRNAs) targeting *JARID2* and the *AAVS1* negative control gRNA (**Fig. S1a**). Peripheral blood (PB) engraftment of human CD45+ (hCD45+) cells reached saturation by eight weeks post-transplantation and remained stable at high levels throughout the study period (**Fig. 1a**). The percentage of targeted cells for the two gRNAs targeting *JARID2* was higher than the *AAVS1* control, but all remained stable over one year (**Fig. 1b**). After one-year post-transplant, there were no significant difference in the peripheral blood lineage differentiation spectrum between the groups (**Fig. 1c**) in terms of proportions of myeloid cells (CD33+), B-cells (CD19+) and T-cells CD3+). Similarly, lineage output was observed in the BM (**Fig. S1b**) and complete blood count (CBC) analysis revealed no significant differences between the groups (**Fig. 1d**; **Fig. S1c**). Histological analysis using hematoxylin and eosin (H&E) and reticulin staining showed no significant pathological changes in the bone marrow or spleens of transplanted mice (**Fig. 1e**). In summary, long-term *JARID2* deficiency in human UCB CD34+ cells did not induce notable hematopoietic pathologies *in vivo*, suggesting that manipulating JARID2 could be a safe translational approach. This is consistent with the observation that *JARID2* is not mutated in human clonal hematopoiesis (CH) or is deleted as a founding event in human blood cancers (*27*), suggesting that *JARID2* loss-of-function cannot operate as an oncogenic event in isolation.

**Fig. 1:**
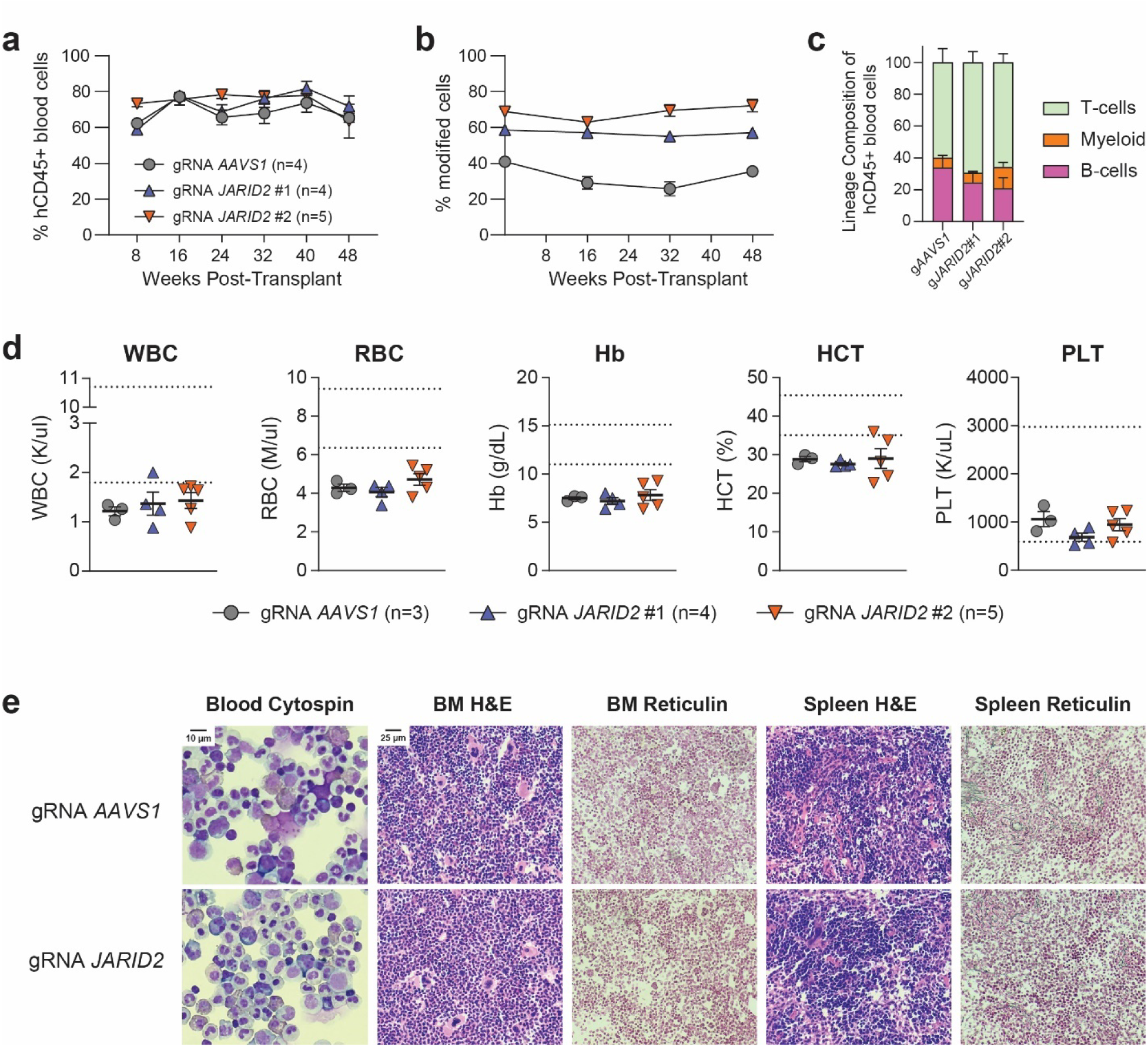
Long-Term Deletion of JARID2 Produces Durable Hematopoiesis Without Transformation. (a) Longitudinal analysis of human CD45+ chimerism in peripheral blood (PB) of NSG mice transplanted with cord blood CD34+ cells targeted with guide RNAs for *AAVS1* (n=4), *JARID2*#1 (n=5) or *JARID2*#2 (n=5). (b) Longitudinal analysis of modified alleles from CRISPR edits in human CD45+ PB cells in transplanted NSG mice. (c) Lineage composition of human CD45+ PB cells one-year post-transplant (B-cells = CD19+, T-cells = CD3+, myeloid cells = CD33+). (d) Complete blood count (CBC) analysis of white blood cells (WBC), red blood cells (RBC), hemoglobin (Hb), hematocrit (HCT) and platelets (PLT) of recipient mice one-year post-transplant. Dashed lines represent normal range. (e) Representative histology of PB smear, bone marrow (BM) cytospins, and spleen (SPL) harvested one-year after transplantation of CRISPR-edited human CD34+ cells into NSG mice. Scale bars - 10µm (PB) and 25µm (BM and SPL).

### JARID2 Inhibition Enhances Functional Output of Human HSPCs in vivo

To facilitate genetic inhibition of *JARID2* in human CD34+ cells, lentiviral vectors expressing short hairpin RNAs (shRNAs) and a GFP reporter were constructed (*28*). The two most effective shRNAs targeting *JARID2* (**Fig. S2a**) relative to the negative control *Renilla* shRNA (shRen) were chosen for experimentation. UCB CD34+ cells were transduced with GFP-expressing lentiviruses and 48-hours post-transduction 4.0×10^4^ GFP+CD34+ cells were transplanted into sublethally irradiated NSG mice. Additionally, excess cells were cultured for eight-days *ex vivo* to observe effects of reprogramming mediated by *JARID2* inhibition. Transduction with *JARID2* shRNAs produced more overall cellular output (**Fig. S2b**), increased numbers of CD34+ cells (**Fig. S2c**), and increased colony forming unit (CFU) activity from a fixed volume of cell culture equivalent (**Fig. S2d**).

Genetic inhibition by *JARID2*-knockdown (*JARID2*-KD) significantly improved peripheral blood engraftment of UCB CD34+ cells relative to the sh*Ren* control group (**Fig. 2a**) without altering lineage differentiation bias (**Fig. 2b**) or overall blood counts (**Fig. S2e**). At 20-weeks post-transplant, *JARID2* inhibition was sustained in targeted BM CD34+ cells (**Fig. S2f**) and engraftment of *JARID2*-KD cells was significantly increased in the BM (**Fig. 2c**). Moreover, flow cytometric analysis (**Fig. S2g**) revealed increased numbers of GFP+ HSPCs including hematopoietic stem cells (HSCs; CD34+ CD38- CD90+ CD45RA-), multipotent progenitors (MPPs; CD34+ CD38- CD90- CD45RA-) and multilymphoid progenitors (MLPs; CD34+ CD38- CD90- CD45RA+) resulting from *JARID2*-KD (**Fig. 2d**). These results show constitutive *JARID2* suppression can increase functionality from transplantation of a limited number of human HSPCs.

**Fig. 2:**
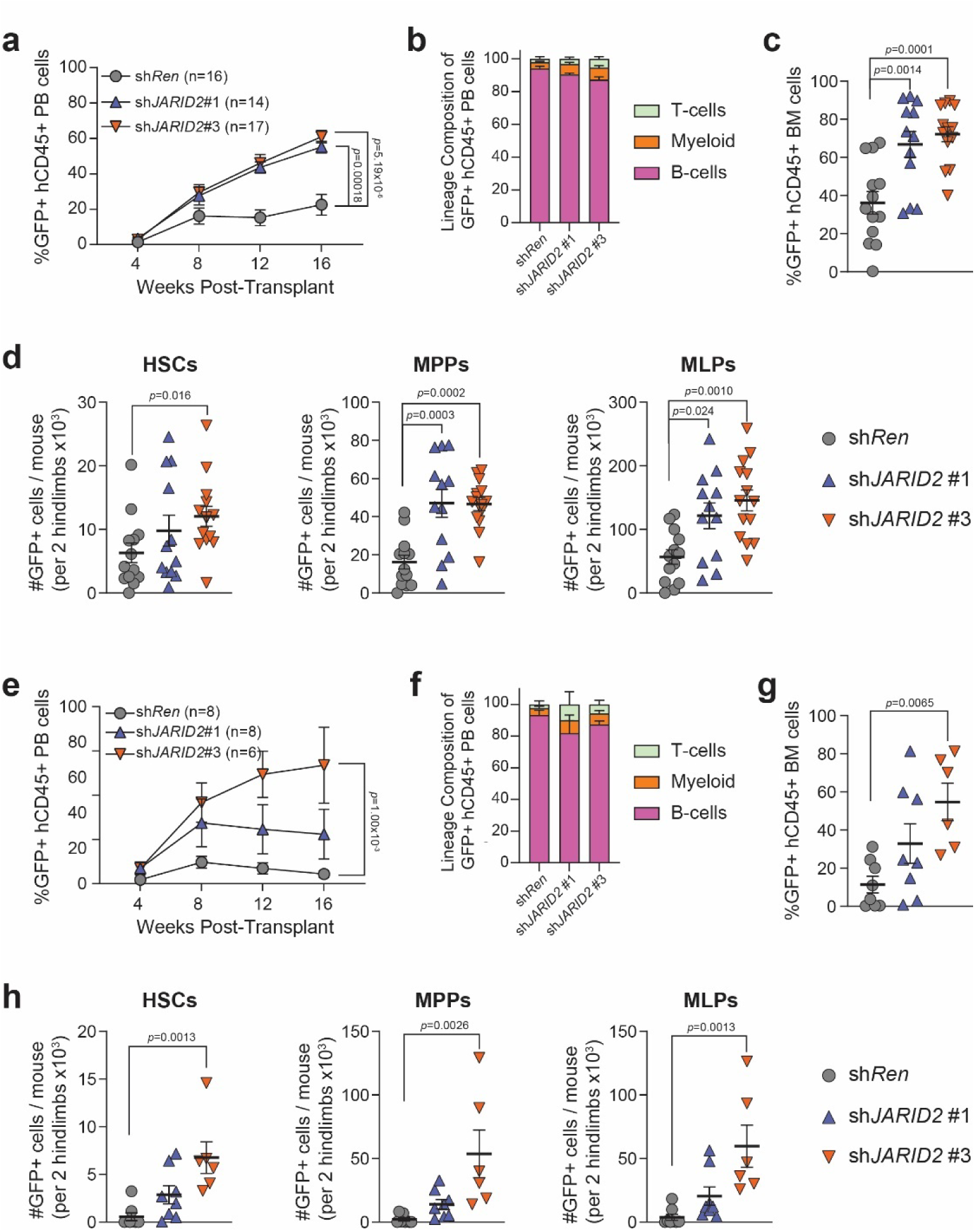
JARID2 Inhibition Enhances Functional Output of Human HSPCs in vivo. (a) Peripheral blood (PB) engraftment of cord blood cells transduced with indicated constitutive lentiviral shRNAs. (b) Lineage composition of GFP+ PB cells 16-weeks post-transplant. (c) Engraftment of GFP+ human cells in the BM of recipient mice 20-weeks post-transplant. (d) Quantification of GFP+ human HSPC populations in the BM of recipient mice 20-weeks post-transplant (sh*Ren* n=16, sh*JARID2*#1 n=14, sh*JARID2*#3 n=17). (e) PB engraftment of cord blood cells transduced with indicated constitutive lentiviral shRNAs in secondary recipients. (f) Lineage composition of GFP+ PB cells 16-weeks post-secondary transplant. (g) Engraftment of GFP+ human cells in the BM of secondary recipient mice 20-weeks post-transplant. (h) Quantification of GFP+ human HSPC populations in the BM of recipient mice 20-weeks post-transplant (sh*Ren* n=8, sh*JARID2*#1 n=6, sh*JARID2*#3 n=6).

To more stringently assess long-term HSC self-renewal, 1.0×10^6^ CD34+ cells from primary recipients were transplanted into secondary NSG mice. *JARID2*-KD cells maintained higher peripheral blood engraftment in secondary recipients (**Fig. 2e**) with normal differentiation (**Fig. 2f**). *JARID2*-KD cells showed increased overall engraftment in the BM of secondary recipients (**Fig. 2g**) as well as increased numbers of HSPC populations (**Fig. 2h**). Notably, sh*JARID2*#3 produced more pronounced functional enhancement than sh*JARID2*#1, which was likely due to dose-dependent *JARID2* inhibition. Cumulatively, these data show *JARID2* inhibition enhances both the lineage reconstitution and self-renewal capacity of human HSPCs, highlighting therapeutic potential for clinical applications.

### JARID2 Inhibition Enhances HSC Function Without Altering Lineage Specification or Gene Expression Programs

We performed spectral antibody immunophenotypic analysis of primary transplant recipients to more comprehensively characterize the spectrum of differentiated cells produced after *JARID2*-KD in UCB CD34+ cells. Representative t-SNE plots (**Fig. 3a**; **Fig. S3a**) and heatmaps (**Fig. S3b**) reflecting cell distributions demonstrated comparable cell populations between *JARID2*-KD and sh*Ren* control groups. In a parallel experiment, transduced UCB CD34+ cells were transplanted into NSGS mice to better assess myeloid output (**Fig. S3c**). No differences in myeloid differentiation potential were observed between the groups (**Fig. 3b**). Collectively, these results further demonstrate that long-term *JARID2* inhibition does not impair multilineage differentiation.

**Fig. 3:**
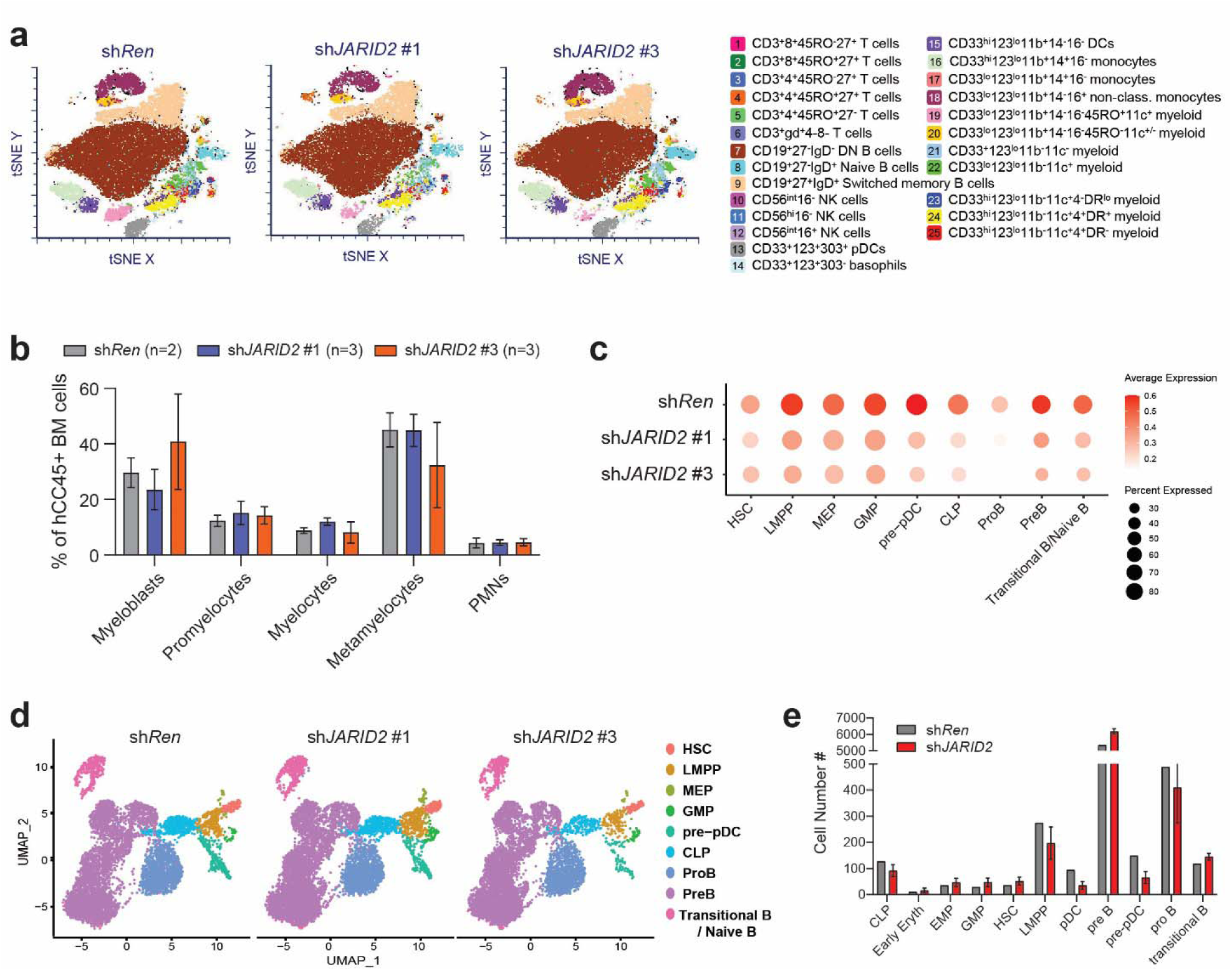
JARID2 Inhibition Enhances HSC Function Without Altering Lineage Specification or Gene Expression Programs. (a) tSNE plots of lineage distribution of human CD45+ cells in BM of recipient mice 20-weeks post-transplant by flow cytometry. (b) Frequency of maturing myeloid cells in NSGS mice transplanted with cord blood cells transduced with indicated constitutive lentiviral shRNAs 20-weeks post-transplant. (c) *JARID2* expression across indicated cell populations showing knockdown efficiency. (d) UMAP of scRNA-seq data for hCD34+ cells isolated 20-weeks post-transplantation. Cell populations are identified by marker genes. (e) Numbers of cells of indicated genotypes per cluster based on Azimuth cell type annotations.

To evaluate stem and progenitor cell composition in more granularity, human CD34+ cells were isolated from the BM of primary NSG recipients and subject to single-cell RNA sequencing. Cell identities were assigned based on selected signature genes (**Fig. S3d**) and unsupervised uniform manifold approximation and projection (UMAP) clustering identified distinct hematopoietic lineages (**Fig. S3e**). Efficient knockdown of *JARID2* was observed across all sh*JARID2* populations (**Fig. 3c**). Quantification of cell numbers within each population between each group (**Fig. 3d**) using Azimuth reference dataset within Seurat showed no significant differences in lineage distribution between groups (**Fig. 3e**)(*29, 30*). These data suggest that *JARID2* inhibition enhances functional capacity of human HSPCs without dramatically altering transcriptional networks or lineage specification programs.

### Transient JARID2 Inhibition Imparts Long-Term But Reversible Functional Benefits to Human HSPCs

We next aimed to more closely simulate clinical stem cell collection and transplantation protocols where UCB CD34+ cells would be transiently exposed to *JARID2* inhibition *ex vivo* prior to transplantation, after which the exposure would be ceased. To facilitate this, doxycycline-inducible lentiviruses were engineered to constitutively express a reverse tetracycline-controlled transactivator (rtTA3) protein from a PGK promoter with separate co-expression of shRNA of interest and blue fluorescent protein (BFP) driven by a Tet-On 3G (T3G) promoter in the presence of doxycycline (**Fig. S4a**). Dose titration experiments identified 200 ng/mL doxycycline as optimal, achieving effective knockdown with minimal cytotoxicity. Vector functionality was validated in HEL cells by Western blot, confirming efficient JARID2 suppression upon doxycycline exposure which was reversed when doxycycline was withdrawn (**Fig. S4b**). The kinetics of vector activity loss after doxycycline withdrawal was confirmed in both the HEL cell line (**Fig. S4c**) and UCB CD34+ cells (**Fig. S4d**). Given that JARID2 is a PRC2 co-factor and prior studies reported 90% of promoters bound by JARID2 in human stem cells were also associated with PCR2 binding (EZH2 / SUZ12)(*31, 32*), we generated *EZH2*-targeting shRNAs in this inducible lentivirus system (**Fig. S4e**) to determine if similar effects would be recapitulated by PRC2 inhibition. This would begin to determine if the mechanism of functional enhancement of human HSPCs following *JARID2* inhibition was PCR2-dependent.

UCB CD34+ cells per well were transduced with inducible vectors carrying shRNAs targeting *Renilla* (negative control), *JARID2* or *EZH2*. After 48-hour exposure to doxycycline, 4.0×10^4^ transduced (BFP+) CD34+ cells were purified and cultured for eight days with doxycycline. The entire progeny of these cultures were transplanted into NSG mice. BFP+ hCD45+ cells became undetectable in the peripheral blood of recipient mice by four-weeks post-transplant, suggesting that shRNA expression was no longer active *in vivo* in the absence of doxycycline (**Fig. S4f**). Even with only transient inhibition *ex vivo*, *JARID2*-KD enhanced the peripheral blood engraftment of UCB CD34+ cells compared to the sh*Ren* control group (**Fig. 4a**) with normal lineage differentiation (**Fig. 4b**). In stark contrast, this effect was not observed from the sh*EZH2* groups (**Fig. 4a**). The divergent effects observed between *JARID2* and *EZH2* inhibition suggest the function of JARID2 in HSCs may not be completely tied to its PRC2 co-factor activity, in contrast to what has been reported in human embryonic stem (ES) cells (*32*). BM engraftment of human cells was more variable but significantly lower in the sh*EZH2* groups (**Fig. 4c**). But human HSPC numbers were consistently increased in the group that received transient inhibition from the more potent *JARID2* shRNA (**Fig. 4d**).

**Fig. 4:**
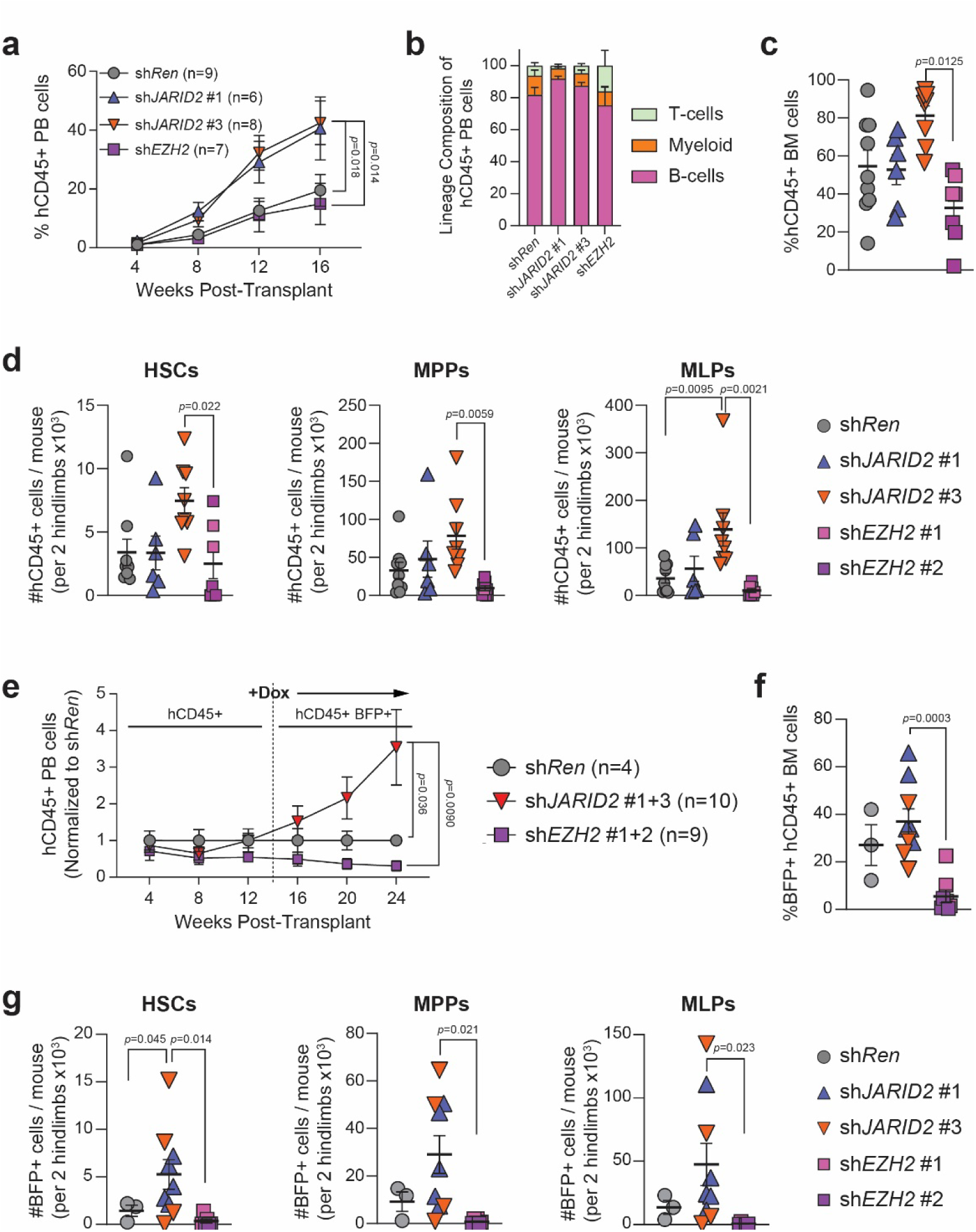
Transient JARID2 Inhibition Imparts Long-Term But Reversible Functional Benefits to Human HSPCs. (a) Peripheral blood (PB) engraftment of cord blood cells transduced with indicated inducible lentiviral shRNAs induced for eight-days *ex vivo* prior to transplant. (b) Lineage composition of hD45+ PB cells 16-weeks post-transplant. (c) Engraftment of hCD45+ cells in the BM of recipient mice 20-weeks post-transplant. (d) Quantification of hCD45+ HSPC populations in the BM of recipient mice 20-weeks post-transplant (sh*Ren* n=9, sh*JARID2*#1 n=6, sh*JARID2*#3 n=8, sh*EZH2* n=7). (e) PB engraftment of cord blood cells transduced with indicated inducible shRNAs induced *in vivo* at 14-weeks post-transplant (normalized to sh*Ren*). (f) Engraftment of BFP+ hCD45+ cells in the BM of recipient mice (shRNAs for *JARID2* and *EZH2* are combined for statistical comparison). (g) Quantification of BFP+ hCD45+ HSPC populations in the BM of recipient mice (sh*Ren* n=3, sh*JARID2*#1+3 n=9, sh*EZH2*#1+2 n=8).

To evaluate long-term HSC self-renewal, 4.0×10^5^ CD34+ cells were isolated from the BM of primary recipients and transplanted into secondary NSG mice. In this setting, PB engraftment of human cells was comparable amongst all groups (**Fig. S4g**). Western blot of human CD34+ cells from the BM of secondary recipients confirmed restoration of JARID2 protein levels following doxycycline withdrawal (**Fig. S4h**), further demonstrating the reversibility of JARID2 system *in vivo*. These data suggest that transient *JARID2* inhibition can provide an immediate benefit to human HSPCs that enhances initial reconstitution of human cells, but this phenotype is lost when JARID2 protein levels are restored in the absence of doxycycline as long-term self-renewal in secondary recipients was comparable to the sh*Ren* control group. Thus, any functional reprogramming resulting from *JARID2* inhibition is not permanent and can be reset *in vivo*.

We leveraged this inducible system to determine if *JARID2* inhibition could directly reprogram human HSPCs *in vivo*. To test this, UCB CD34+ cells were transduced with inducible lentiviral shRNAs in the absence of doxycycline and transplanted into NSG mice. In the absence of induction, peripheral blood engraftment of human cells was comparable amongst all groups. 14-weeks post-transplant, expression of shRNAs was induced through administration of doxycycline in rodent chow. Immediately following induction, sh*JARID2* groups displayed a significant increase in peripheral blood engraftment relative to the sh*Ren* control group, whereas the sh*EZH2* group showed a decline in engraftment levels (**Fig. 4e**). After 12-weeks of doxycycline induction, there was increased engraftment of human cells in the BM (**Fig. 4f**) and increased numbers of BFP+ HSPCs (**Fig. 4g**) from the *JARID2*-KD groups. This experiment shows human HSPCs can be directly reprogrammed *in vivo* by *JARID2* inhibition to enhance functional output.

### Cell Culture Changes the Phenotypic Identify of Functional Human Repopulating Cells

After *ex vivo* culture, the CD34+ cell population contains a mixture of HSPCs, most of which lack long-term repopulating potential. Our prior data in mouse models showed that genetic inhibition of *Jarid2* conveyed long-term repopulating potential to MPPs (*22*), a cell population that possesses only transient engraftment potential under normal circumstances. It is possible a similar mechanism operates in human cells and the enhanced functional output of UCB CD34+ cells following *JARID2*-KD is due to differential effects on specific HSPC subpopulations. To examine this, UCB CD34+ cells were transduced with GFP-expression lentiviral vectors constitutively expressing shRNAs targeting *JARID2* or *Renilla* negative control. In this case, following *ex vivo* culture 1250 HSCs (CD34+ CD38- CD90+ CD45RA-) or 1500 MPPs (CD34+ CD38- CD90- CD45RA-) were purified based on canonical human cell surface markers and transplanted into NSG mice. At 12-weeks post-transplant, peripheral blood engraftment of HSCs receiving *JARID2*-KD exhibited higher engraftment than the sh*Ren* control group (**Fig. S5a**). However, MPPs from all groups failed to engraft (**Fig. S5b**) and overall engraftment levels were markedly lower than that observed in prior experiments. This suggested that the immunophenotype used to identify native human HSPC population may be influenced by *ex vivo* culture and/or lentiviral transduction.

Colony forming unit (CFU) assays were performed to define the phenotype of functional HSCs after *ex vivo* culture. While used as a marker of long-term human HSCs *in vivo* (*33–35*), CD49f was no longer a reliable marker of functional HSCs after *ex vivo* culture as all CD49f+ cells failed to generate colonies (**Fig. S5c**). Rather, the CD34+EPCR+CD90+CD49f-phenotype enriched for most robust CFU activity (**Fig. S5c,d**). Eight days post-transduction, *JARID2*-KD increased the number of CD34+EPCR+CD90+CD49f-cells in culture (**Fig. S5e**) and enhanced CFU activity from this population on a per cell basis, which was enhanced in replating experiments (**Fig. S5f**). CITE-seq was performed to integrate single cell transcriptomics with immunophenotype of cells after *ex vivo* culture (**Fig. S5g**). As CD49f+ cells lacked CFU potential after culture, this population (**Fig. S5h**) was excluded from the final UMAP (**Fig. 5a**). HSCs were identified by specific expression of canonical human HSC-defining genes (**Fig. S5i**). However, in addition to a cluster of cells expressing well-described human HSC-defining genes such as *HLF*, expression of other HSC-specific genes such as *MECOM* and *MEIS1* spread to another cluster of cells after *ex vivo* culture (**Fig. 5b**). This emergent population was expanded after *JARID2*-KD and also marked by cell surface expression of EPCR (CD201) and CD90 (**Fig. 5c**). This emergent population was annotated as megakaryocyte-erythroid progenitors (MEPs) based on transcriptional signature (**Fig. 5a**), with expression of CD71 demarcating this population from HSCs (**Fig. 5c**). However, CD71 has also been noted as a marker of activated HSCs (*36*) and given the ectopic expression of some HSC-specific genes in this population and it’s expansion after *JARID2*-KD, we hypothesized this may represent the cell population that is functionally augmented after *JARID2* inhibition.

**Fig. 5:**
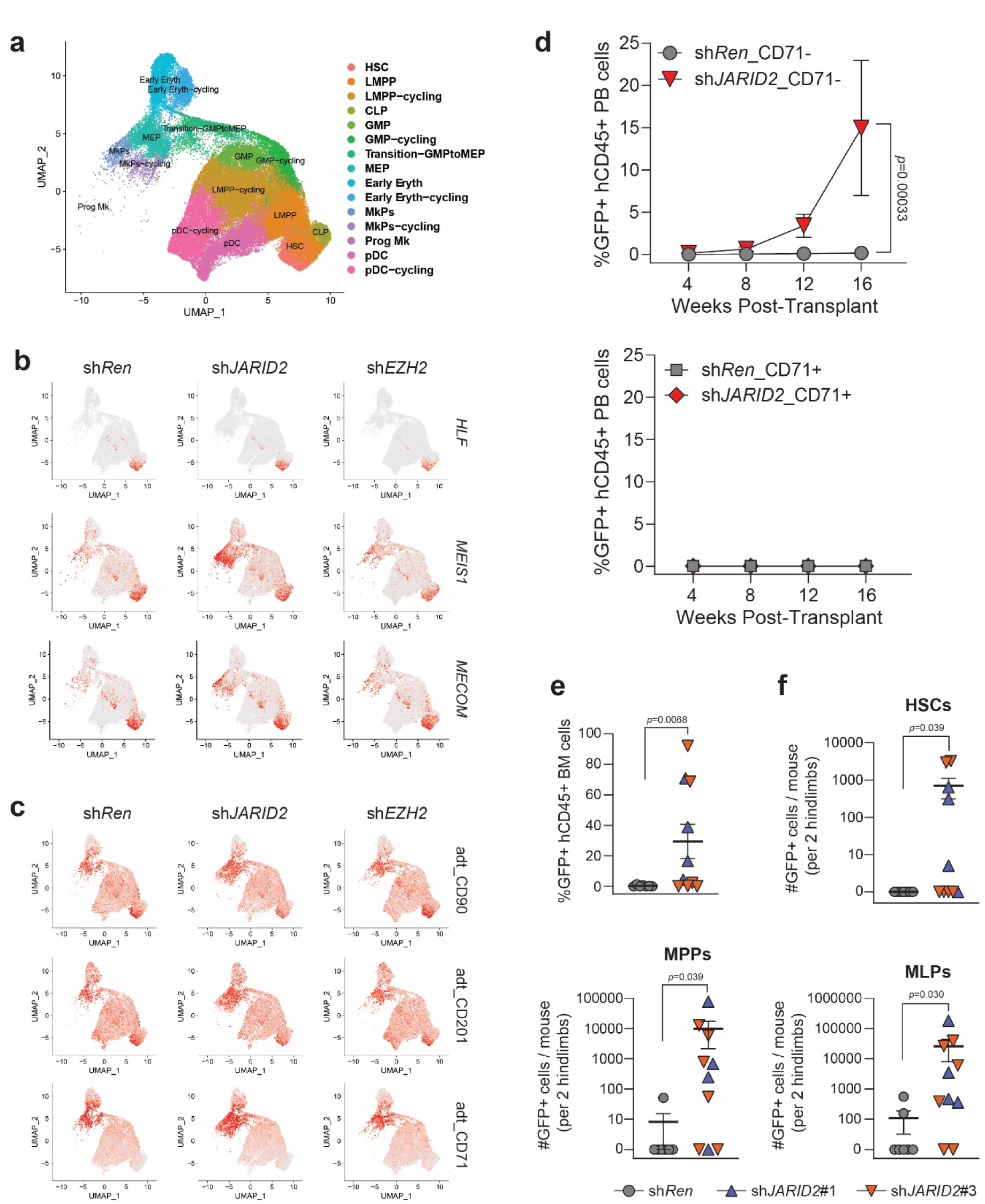
Cell Culture Changes the Phenotypic Identify of Functional Human Repopulating Cells. (a) UMAP of CITE-seq data from GFP+CD34+ cells after eight-days culture (CD49f+ cells excluded). (b) Feature plots showing expression of HSC marker genes *HLF*, *MEIS1* and *MECOM*. (c) Feature plots showing expression of antibody-derived tags (adt) for surface proteins CD90, CD201 (EPCR) and CD71. (d) PB engraftment in recipient mice transplanted with 1000 cells of indicated HSC (GFP+CD34+CD90+ EPCR+ CD49f-) phenotype (CD71+ / CD71-) transduced with indicated lentiviral shRNAs. (e) Engraftment of GFP+ human cells in the BM of recipient mice 20-weeks post-transplant.

To analyze the functional potential of this emergent population after *JARID2*-KD, UCB CD34+ cells were transduced with constitutive lentiviral shRNA vectors and after eight days of culture, equal numbers of CD34+EPCR+CD90+CD49f-cells were purified from each group with additional separation based on CD71 expression (**Fig. S5j**) and transplanted into NSG mice. Only the CD71-cell fraction displayed long-term engraftment potential which was enhanced by *JARID2*-KD (**Fig. 5d**). *JARID2*-KD in CD34+EPCR+CD90+CD49f-CD71-cells also resulted in higher human cell chimerism in the BM of recipient mice (**Fig. 5e**) and increased numbers of HSPCs (**Fig. 5f**). Similar results were observed from serial CFU replating of these presumptive HSC populations (**Fig. S5k**). Although *JARID2*-KD expanded a population of cells with HSC transcriptional and phenotypic characteristics in culture, ultimately this CD71+ cell fraction possessed no reconstitution capacity *in vivo*. Our data reveal that functional repopulating activity was contained within the CD34+EPCR+CD90+CD49f-CD71-cell fraction after culture manipulation.

### JARID2 Knockdown Produces Subtle Molecular Effects in HSPCs that are Distinct From PRC2 Inhibition

To further investigate the molecular mechanisms underlying the enhanced functionality of HSPCs upon *JARID2* inhibition, epigenetic profiles of UCB CD34+ cells following *JARID2* and *EZH2* knockdown were compared. Western blot analysis after *ex vivo* culture confirmed that *EZH2* knockdown led to significant depletion of H3K27me3 in HSPCs, consistent with its central role in PRC2-mediated histone methylation (**Fig. 6a**). In contrast, *JARID2* knockdown did not significantly alter global H3K27me3 levels (**Fig. 6a**), suggesting a distinct role from the PRC2 core complex in human HSPCs. To assess if specific loci were affected, CUT&RUN and CUT&TAG was performed for a variety of epigenetic modifications on purified GFP+CD34+EPCR+CD90+CD49f-CD71-cells after eight days of culture. Experiments were performed on three independent UCB samples as biological replicates and compared to the freshly isolated UCB CD34+ cells as a reference control. Consistent with Western blot results, H3K27me3 peaks were largely maintained in *JARID2*-KD samples, but markedly reduced in *EZH2*-KD samples (**Fig. 6b,c**). Despite these epigenetic distinctions, the inherent biological variation of individual UCB donors was the strongest driver of sample variability (**Fig. 6d**). After removing the effects of individual biological replicates, pathway analysis of H3K27me3-enriched regions (**Table S1**) did not identify significantly enriched terms (**Fig. S6a**). This effect was consistent across multiple epigenetic marks profiled such as H3K4me3, H2AK119ub and CTCF (**Fig. S6b**). Transcriptional profiling by bulk RNA-seq of the same cell populations revealed a similar effect (**Fig. 6e**). The variability of genetic background between different donors resulted in identification of very few differentially expressed genes (DEGs) between comparisons (**Table S2**), none of which seemed to explain the observed functional results (**Fig. S6c**).

**Fig. 6:**
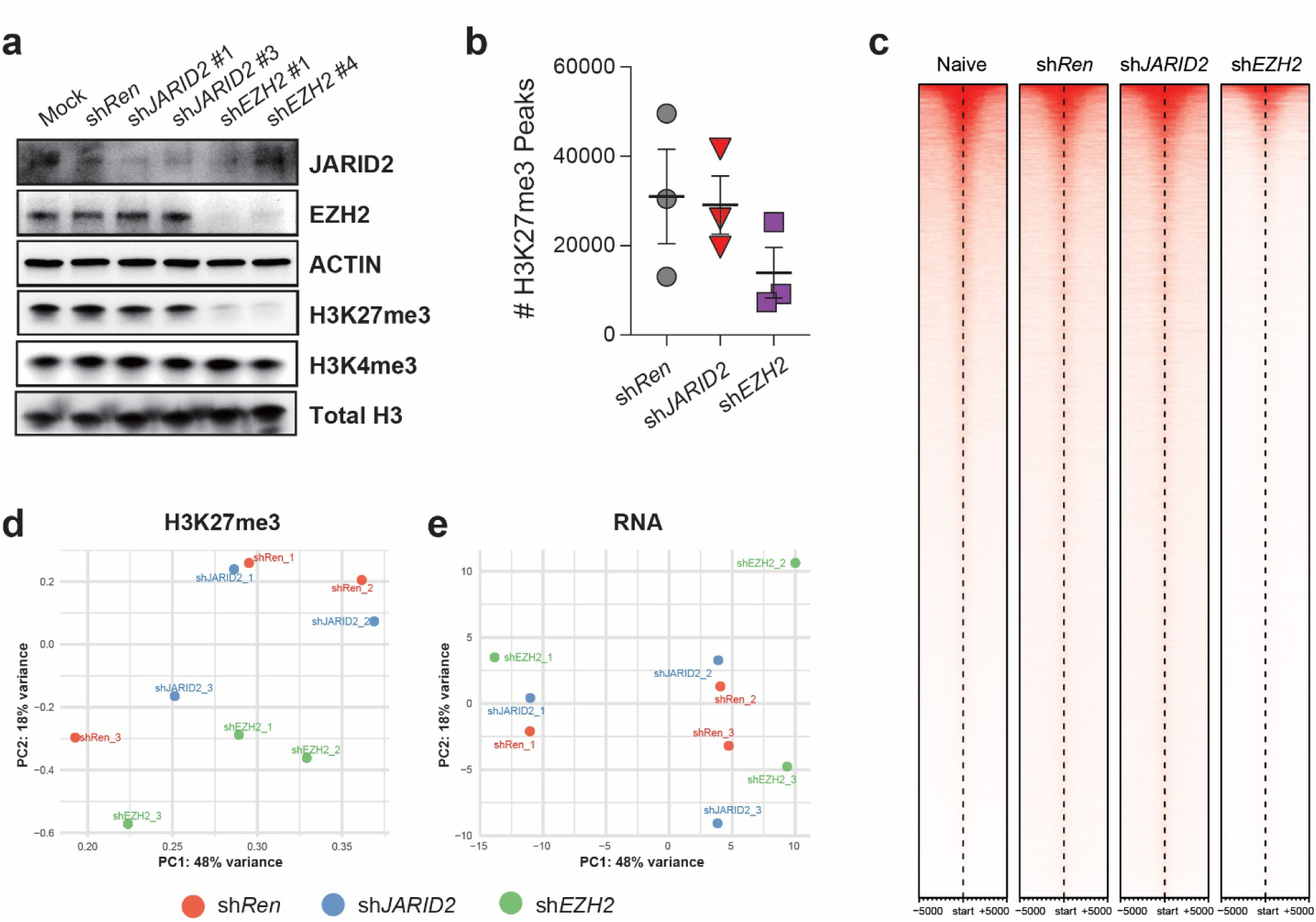
JARID2 Knockdown Produces Subtle Molecular Effects in HSPCs that are Distinct From PRC2 Inhibition. (a) Western blot analysis of cord blood CD34+ cells transduced with indicated shRNAs after eight-days of *ex vivo* culture. (b) H3K27me3 peak counts from CUT&Tag analysis of GFP+CD49f-CD90+CD34+EPCR+CD71- cells transduced with indicated shRNAs after eight-days of *ex vivo* culture. (c) Heatmaps of H3K27me3 signal intensity centered at transcription start sites (±5 kb) for genes identified in all four experimental conditions. Data represent combined results from three biological replicates per condition. Heatmaps are sorted by decreasing H3K27me3 signal intensity in naïve UCB CD34+ reference sample. (d) Principal component analysis (PCA) of H3K27me3 peak profiles from CUT&Tag analysis of GFP+CD49f-CD90+CD34+EPCR+CD71- cells transduced with indicated shRNAs after eight-days of *ex vivo* culture. (e) PCA of RNA-seq gene expression profiling of GFP+CD49f-CD90+CD34+EPCR+CD71- cells transduced with indicated shRNAs after eight-days of *ex vivo* culture.

### JARID2 Knockdown Reinforces Stem Cell Gene Expression Programs and Reprograms MPPs in vivo

As cell culture conditions modified transcriptomic and immunophenotypic properties of HSCs, to uncover true effects of *JARID2*-KD in HSPCs we revisited scRNA-seq data generated from GFP+CD34+ cells isolated from recipient mice after transplant, reasoning this would provide sufficient time to re-establish native cell surface markers. Within the annotated HSC cell cluster (**Fig. 7a**), *JARID2*-KD increased expression of well-annotated HSC-defining genes (*37, 38*) such as *HLF*, *MLLT3*, *MYCT1*, and *MEIS1* relative to control sh*Ren* HSCs (**Fig. 7b**). Due to the limited dataset size, only a few individual genes reached statistical significance across conditions. To address this, we incorporated a curated list of HSC-defining genes from a recent study to compute a HSC module score (*38*). This HSC module score was significantly elevated in sh*JARID2* HSCs relative to control sh*Ren* HSCs (**Fig. 7c**). Looking across other cell populations, while as expected the HSC module score peaked in the HSC cluster, elevated scores were also detected in other *JARID2*-KD HSPC populations (**Fig. S7a**), suggesting *JARID2* inhibition may reprogram some of these populations with HSC-like potential *in vivo*. To test this, HSCs, MPPs and MLPs were isolated from the BM of secondary recipients of constitutive shRNA transduced UCB CD34+ cells 20-weeks post-transplant and subject to *in vitro* functional analysis via CFU assay. *JARID2*-KD significantly enhanced serial replating of both HSC and MPP cell fractions, but not MLPs (**Fig. 7d; Fig. S7b**). Thus, in human HSPCs, *JARID2* inhibition not only enhances function of HSCs, but also conveys HSC-like potential to the MPPs which lacks self-renewal capacity under normal conditions, analogous to our prior results from murine models (*22*).

**Fig. 7:**
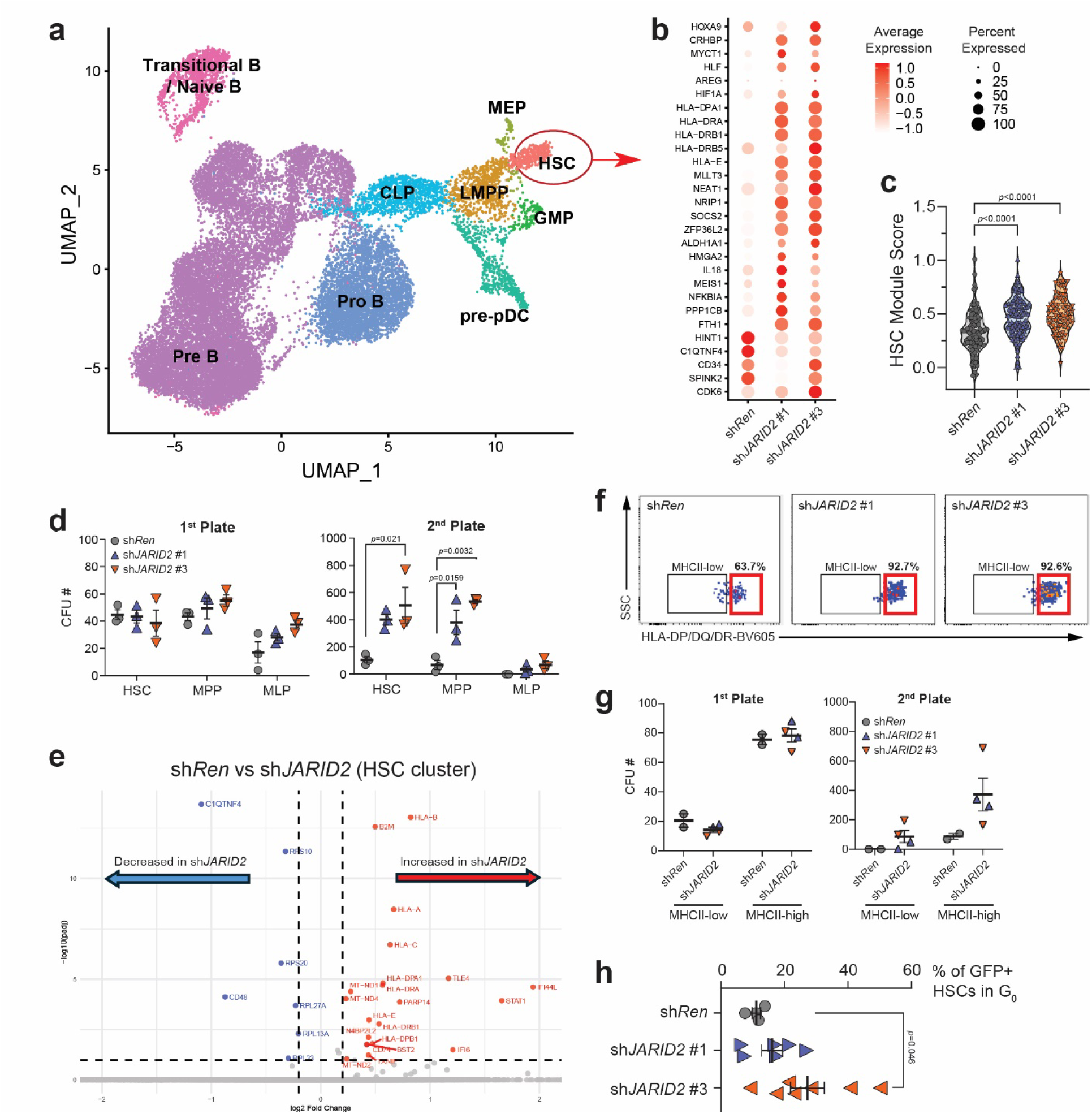
JARID2 Knockdown Reinforces Stem Cell Gene Expression Programs and Reprograms MPPs in vivo. (a) UMAP of scRNA-seq data for hCD34+ cells sorted from NSG mice transplanted with cord blood cells transduced with constitutive lentiviral shRNAs 20-weeks post-transplant. Cell populations are identified by marker genes. sh*Ren*, sh*JARID2*#1 and sh*JARID2*#3 samples are combined, the HSC population is highlighted. (b) Dot plot highlighting differential expression of a curated HSC signature gene set (Aguadé-Gorgorió et al., 2024, Nature) amongst shRNA genotypes within the HSC population. (c) HSC module scores calculated using the Seurat package based on the curated HSC signature gene list. (d) Serial replating showing CFU efficiency of indicated HSPC populations isolated from BM of recipient mice 20-weeks post-transplant. (e) Volcano plot showing differentially expressed genes between sh*JARID2* and sh*Ren* cells within the HSC population. (f) Representative flow cytometry plots showing the proportion of MHCII-high (red boxes) HSCs from each genotype from the BM of recipient mice 20-weeks post-transplant. (g) Serial replating showing CFU efficiency of indicated HSPC genotypes separated by MHCII levels isolated from BM of recipient mice 20-weeks post-transplant. (h) Cell cycle analysis by flow cytometry showing proportion of quiescent (G_0_) HSCs of each genotype from the BM of recipient mice 20-weeks post-transplant.

To investigate the molecular mechanisms underlying the enhanced function observed following *JARID2* inhibition, differential gene expression analysis was compared specifically within the HSC population. While most downregulated genes were related to ribosome biogenesis (**Fig. S7c**), the most significantly upregulated genes in *JARID2*-KD HSCs were largely related to the major histocompatibility complex class II (MHCII; **Fig. 7e**). Recent studies have suggested a role for MHCII in HSC regulation through antigen presentation as a mechanism of safeguarding the integrity of the HSC pool (*39, 40*). Flow cytometry validated the scRNA-seq data and confirmed higher expression of MHCII proteins on the surface of *JARID2*-KD HSCs (**Fig. 7f**). As MHCII has been suggested to mark long-term HSCs in mice (*39*), to assess if increased MHCII marked more functional human cells after *JARID2*-KD, HSCs from recipient mice transplanted with UCB CD34+ cells transduced with constitutive shRNA vectors were sorted based on MHCII expression and subject to CFU assay. Serial replating showed that *JARID2*-KD enhanced functionality of both MHCII-negative and MHCII-high HSCs, but most of the CFU potential was contained in the MHCII-high fraction which was dramatically enhanced following *JARID2* inhibition (**Fig. 7g**). In both mice and human, MHCII-high HSCs show reduced cycling and apoptosis and are resistant to stress-induced proliferation, linked to MHCII-high HSCs residing in a deeper quiescent state (*39, 40*). We performed cell cycle analysis of post-transplant transduced HSCs (**Fig. S7d**) which confirmed that *JARID2*-KD HSCs contained a significantly higher proportion of cells in the quiescent G_0_ phase (**Fig. 7h**). As deeper quiescence is a hallmark of more functional HSCs, this presents one potential mechanism through which *JARID2* inhibition can enhance the long-term reconstitution of human HSCs.

### JARID2 Inhibition Upregulates STAT1 to Enhance Function of Human HSPCs

A recent study showed STAT1 is essential for the maintenance of MHCII-high HSCs (*40*). In addition to MCHII components, *STAT1* was one of the most significantly upregulated genes in *JARID2*-KD HSCs (**Fig. 7e**; **Fig. 8a**). From this data, scatterplot analysis using Seurat revealed a correlation between cells with the highest *STAT1* expression levels in *JARID2*-KD HSCs also having the highest expression of MHCII (**Fig. 8b**) and western blot analysis confirmed upregulation of STAT1 protein in *JARID2*-KD CD34+ cells (**Fig. 8c**). We hypothesized if the mechanism of augmented HSPC function resulting from *JARID2* inhibition was through STAT1 upregulation, that inhibiting *STAT1* in *JARID2*-KD cells may ameliorate their functional enhancement. shRNAs were designed and validated to target *STAT1* and cloned into a BFP-expressing lentiviral vector (**Fig. S8a**). GFP+ CD34+ cells from primary transplant recipients of constitutive shRNA lentiviruses were transduced with *STAT1* shRNA lentiviruses and subject to functional assessment by CFU assay. Results showed that *STAT1* inhibition was able to rescue the increased CFU activity of *JARID2*-KD HSPCs (**Fig. 8d**). While there were no obvious epigenetic changes at the *STAT1* locus in *JARID2*-KD HSCs that would explain the increased expression (**Fig. S8b,c**), collectively these findings suggest that *JARID2* inhibition enhances human HSPC functionality at least in part by upregulating *STAT1* and MHCII while promoting a quiescent gene expression signature characteristic of long-term self-renewing HSCs.

**Fig. 8:**
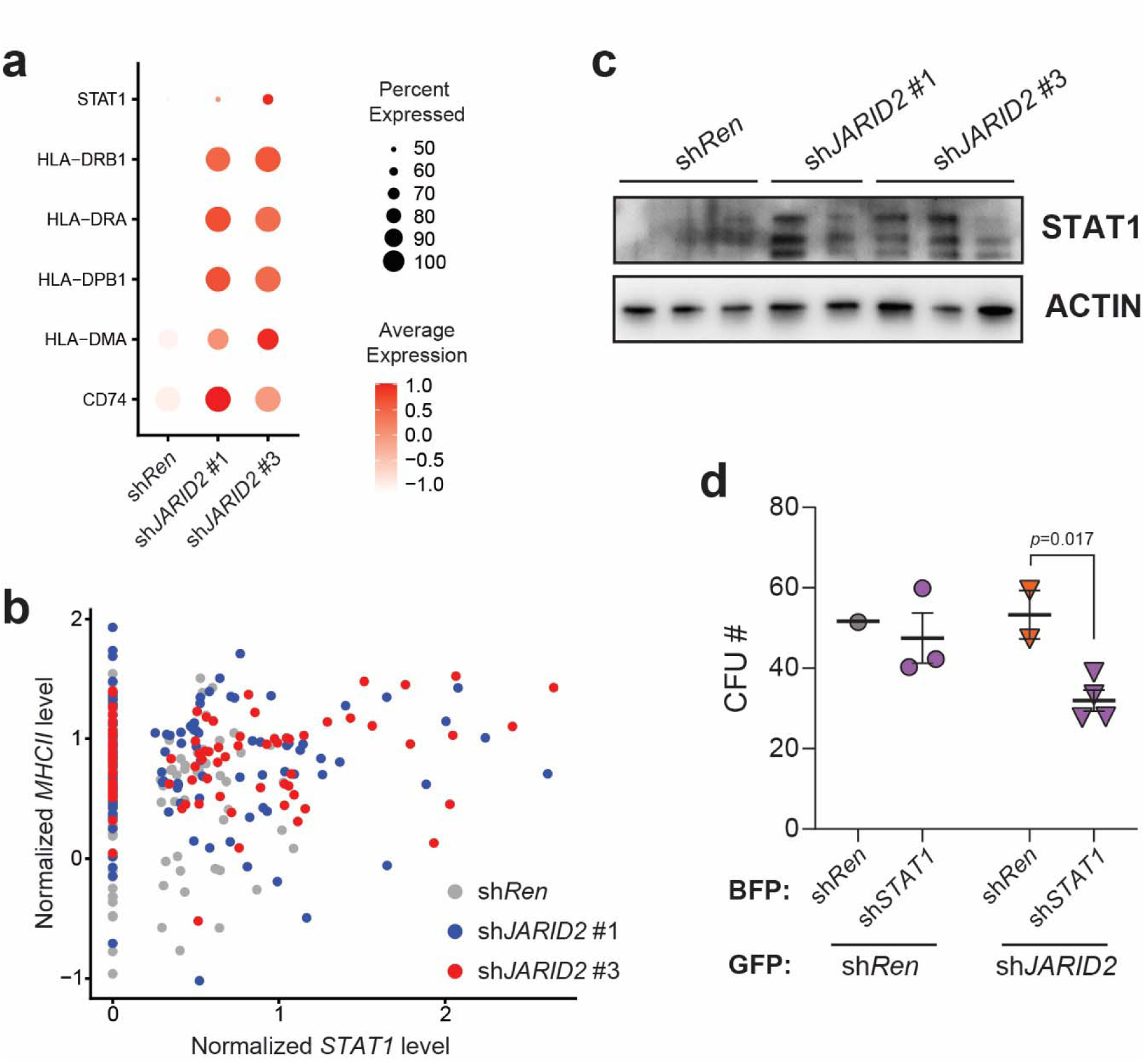
JARID2 Inhibition Upregulates STAT1 to Enhance Function of Human HSPCs. (a) Dot plot showing expression of *STAT1* and *MHCII* components in HSCs of indicated genotypes derived from scRNA-seq data. (b) Scatterplot showing the correlation between *STAT1* mRNA levels and MHCII gene module scores in individual HSCs. Each dot represents a single cell within the defined HSC population. (c) Western blot showing STAT1 protein levels in hCD34+ cells isolated from recipient mice of cord blood cells transduced with indicated lentiviral shRNAs 20-weeks post-transplant. (d) CFU analysis of indicated HSC genotypes isolated from primary recipient mice transduced with indicated BFP+ lentiviral shRNAs.

## DISCUSSION

The ability to effectively propagate functional human HSCs *ex vivo* would have tremendous benefit for BMT and also open new avenues for autologous gene therapy / gene editing applications for human blood disorders. Efforts to propagate transplantable human HSCs *ex vivo* have largely failed due to the repression of self-renewal molecular programs resulting from exposure to cytokines and cell division outside the bone marrow niche. The approach we undertook to tackle this problem was to use acute epigenetic manipulation during *ex vivo* expansion culture to prevent repression of self-renewal genetic networks. Each step of hematopoietic differentiation is associated with modulation of the epigenome as new cell fates are adopted. As epigenetic modifications are reversible and dynamically remodeled during normal development, we reasoned manipulation of the epigenetic machinery could provide a novel approach to biasing HSC fate decisions towards self-renewal and expand functional HSCs. Prior work from our lab identified a crucial role for the PRC2 co-factor Jarid2 in epigenetic repression of self-renewal pathways in developmental hematopoiesis (*22*). In mice, Jarid2 is responsible for recruiting PRC2 to epigenetically repress self-renewal genes in MPPs during HSC differentiation via establishment of the repressive H3K27me3. The goal of this study was to inhibit this function of JARID2 during hematopoietic fate decisions during *ex vivo* culture of UCB CD34+ cells to circumvent epigenetic silencing of self-renewal pathways as a mechanism to expand transplantable human cells while maintaining self-renewal potential. Our data show that *JARID2* inhibition enhances output of human UCB cells through multiple mechanisms. In human HSCs, *JARID2* inhibition promotes a quiescent gene expression signature characteristic of long-term self-renewing HSCs, at least partly driven by upregulation of *STAT1* and marked by increased MHCII expression. Inhibition of *JARID2* also appears to reprogram human MPPs *in vivo* to endow this population with some measure of self-renewal potential. This finding is analogous to prior murine studies (*22*) and extends previous observations regarding the role of *JARID2* in human hematopoiesis (*23*). Importantly for potential translational approaches, transient *JARID2* inhibition either through *ex vivo* or *in vivo* inducible shRNA expression, was sufficient to reprogram human HSPCs with enhanced functional potential. This effect was reversible in secondary transplant experiments showing that transient *JARID2* inhibition does not permanently reprogram human HSPCs, which could alleviate concerns about future malignant potential while providing an initial boost of blood count recovery short-term post-transplant.

The observed mechanism of functional enhancement resulting from *JARID2* inhibition in human HSPCs appears to be at least partly attributed to upregulation of *STAT1*. This aligns with recent data from mouse models that show HSCs from *STAT1*-deficient mice are functionally impaired in competitive transplantation assays and *STAT1* is required to maintain protective transcriptional programs in homeostatic HSCs, including inhibition of cell cycling(*40*). In mice, *STAT1* is also required to maintain MHCII-high HSCs as this population is lost upon genetic deletion of *STAT1*(*40*). Although epigenomic analysis did not identify conclusive insight into how *JARID2* inhibition leads to *STAT1* upregulation, this appears to be a mechanism that augments HSC function that is conserved between mouse and human. Another major caveat of this study is that the exact identity of the human HSPCs that obtain functional benefit from *JARID2* inhibition could not be definitively identified because the nature of *ex vivo* culture conditions alters the immunphenotypic profile of repopulating cells. Nevertheless, results from these studies position *JARID2* inhibition as a novel method to enhance human HSPC function from limiting donor stem cell sources which could be implemented to improve clinical BMT applications.

## MATERIALS AND METHODS

### Study Design

This study was designed to determine if genetic inhibition of *JARID2* was sufficient to enhance the functionality of human HSPCs. This could broaden the clinical utility of UCB as a donor source for stem cell transplantation to treat a variety of blood disorders. UCB CD34+ cells were transduced with lentiviruses expressing constitutive or inducible shRNAs targeting *JARID2* and subject to *in vitro* and in *vivo* functional assays. Sample sizes were not predetermined by a statistical method but informed by historical studies from our laboratory. Investigators were not blinded to sample identity and no data were excluded.

### Animals

All animal procedures were approved by the Institutional Animal Care and Use Committee at Washington University. NOD-scid IL2Rgamma^null^ (NSG) mice were obtained from The Jackson Laboratory (#005557). Male and female mice aged 8-12 weeks were used as recipients and were transplanted via retro-orbital injection after 200 rads of gamma irradiation (Cesium-137). Equal numbers of male and female mice were included in the experiments with no gender bias observed.

### Human Cord Blood Samples

Anonymous cord blood specimens were obtained from the St. Louis Cord Blood Bank, St. Louis Barnes-Jewish Hospital, and the Cleveland Cord Blood Center. All cord blood samples were collected and transported with written consent, in accordance with a protocol approved by the Washington University Human Studies Committee and following the Declaration of Helsinki. Mononuclear cells were isolated from the cord blood samples using Ficoll gradient centrifugation. CD34+ cells were then enriched by magnetic selection (Miltenyi Biotec, #130-100-453) and cultured in SFEMII medium (StemCell Technologies, #09605) supplemented with Penicillin-Streptomycin (50 U/mL), human SCF (100 ng/mL), human TPO (100 ng/mL), and human FLT3L (100 ng/mL).

### Cell Sorting and Flow Cytometry

All cells for flow cytometry were stained for >20 minutes at 4°C in Hank’s Balanced Salt Solution (HBSS; Corning, #21021CV) supplemented with Penicillin-Streptomycin (100 U/mL; Fisher Scientific, #MT30002CI), HEPES (10 μM; Life Technologies, #15630080), and fetal bovine serum (2%; Sigma, #14009C). Antibodies were diluted at 1:100. The following fluorophore-conjugated antibodies were used: anti-human CD11b (BUV737, BD Biosciences, #612800; FITC, BioLegend, #101205), CD11c (PE-Cy7, BD Biosciences, #561356), CD123 (BV510, BioLegend, #306022), CD14 (APC-Cy7, BioLegend, #367108; BV650, BioLegend, #301836), CD15 (BV421, BioLegend, #323039; FITC, BD Biosciences, #555401), CD16 (PE, BD Biosciences, #555407; R718, BD Biosciences, #566969), CD19 (Biotin, BioLegend, #328106; BV605, BioLegend, #302244; PE-Cy7, BioLegend, #302215), CD201/EPCR (BV421, BD Biosciences, #743552), CD27 (BV421, BioLegend, #356418), CD3 (AF488, BD Biosciences, #557694; PE, Invitrogen, #12-0038-42), CD33 (APC, BD Biosciences, #561817; BV421, BD Horizon, #565949; FITC, BioLegend, #366620; PE-CF594, BD Biosciences, #562492), CD34 (APC, BioLegend, #343608; Biotin, BioLegend, #343524; PE, BioLegend, #343606), CD38 (APC, BioLegend, #356606; BV605, BioLegend, #356642), CD4 (BV711, BD Biosciences, #563028), CD45 (APC, BioLegend, #368512; BUV395, BD Biosciences, #563792; Biotin, BioLegend, #304004), CD45RA (APC, BioLegend, #304112; APC-Cy7, BioLegend, #304128; BV421, BioLegend, #304130; FITC, BioLegend, #304106; PerCP/Cy5.5, BioLegend, #304121), CD45RO (PacBlue, BioLegend, #304215), CD49f (APC, BioLegend, #313616; APC-Cy7, BioLegend, #313628; Biotin, BioLegend, #313604), CD56 (PE, BD Biosciences, #340363), CD71 (APC, BioLegend, #334108; FITC, BioLegend, #334103), CD8 (BUV496, BD Biosciences, #612942), CD90 (APC, BioLegend, #328114; PE-Cy7, BioLegend, #328124), CD303 (APC-Vio770, Miltenyi Biotec, #130-113-652), γδ TCR (BB700, BD Biosciences, #745944), HLA-DR (BV786, BD Biosciences, #564041), HLA-DR, DP, DQ (BV605, BD Biosciences, #740408), IgD (APC, BD Biosciences, #561303), and Ki67 (PE, BioLegend, #350503; PE-Cy7, BioLegend, #350525). Mouse-specific antibodies included anti-mouse CD45 (BV605, BioLegend, #103140; PE-Cy5, BioLegend, #103110). Streptavidin-BV605 (BioLegend, #405229) was used for detection of biotin-conjugated antibodies. For cell viability staining, Live/Dead Blue (Thermo Fisher, #L23105) or 7-AAD (BioLegend, #420404) was added at 1:100 dilution. After antibody staining, cells were washed and resuspended in HBSS+ containing 7-AAD for cell viability staining prior to flow cytometry analysis. Flow cytometry was performed using either an Attune NxT Flow Cytometer or a Yeti Flow Analyzer provided by the Siteman Flow Cytometry Core. Cell sorting was carried out on a Beckman Coulter MoFlo or BD Aria Ilu SORP sorter.

### Western Blot

Protein samples were separated on pre-cast 4%–15% gradient SDS-PAGE gels (Bio-Rad, #456– 1084) and transferred to nitrocellulose membranes (Millipore, #IPVH00010). Membranes were blocked with 5% BSA and incubated with primary antibodies targeting specific proteins of interest. The following primary antibodies were used: anti-β-actin (Santa Cruz, #sc-47778), EZH2 (Abcam, #ab3748; CST, #5246S), histone H3 (CST, #4499S), H3K27me3 (CST, #9733S), H3K4me3 (CST, #9751S; EpiCypher, #13-0060), HDAC2 (CST, #2540S), JARID2 (Asp1114) (CST, #13594S), and STAT1 (CST, #9172S). After washing with TBS-T, membranes were incubated with HRP-conjugated secondary antibodies specific to either mouse or rabbit IgG, and detection was carried out using a chemiluminescent HRP substrate (Millipore, #WBKLS0100). Imaging was performed using a Bio-Rad imaging system

### CRISPR/Cas9 mediated gene knock out

We designed multiple small guide RNAs (gRNAs) targeting exon 3 of the human *JARID2* gene using the CRISPR design tool available via the UCSC Genome Browser (https://genome.ucsc.edu/). Synthetic gRNAs were synthesized by Synthego and included two sequences targeting exon 3 of *JARID2*: CTTGGTAATGACCAGTCTAA (anti-*JARID2* #1) and GTTGCTAGTGGAGGACACTT (anti-*JARID2* #2). A gRNA targeting the *AAVS1* locus (GGGGCCACTAGGGACAGGAT) was used as a negative control. CD34+ cells were pre-cultured for 12–24 hours before nucleofection with Cas9 ribonucleoprotein (RNP) complexes (IDT, #1074181) assembled with these synthetic gRNAs, following previously published protocols(*41*). Twenty-four hours post-nucleofection, approximately 250,000 cells were transplanted into sublethally irradiated (200 rads) NSGS mice via X-ray-guided intra-tibial injection. A small aliquot of cells was retained for PCR amplicon-based sequencing to assess the initial CRISPR/Cas9 editing efficiency.

### Immunohistochemistry

Mouse tibias were fixed in 10% neutral buffered formalin (Fisher Scientific, #SF100-4) overnight at 4°C. Decalcification was performed using 14% EDTA for 14 days at 4°C. Following fixation and decalcification, tibia samples were submitted to the Washington University Musculoskeletal Histology and Morphometry Core for hematoxylin and eosin (H&E) and reticulin staining. Mouse spleens were collected, fixed in 10% neutral buffered formalin (Fisher Scientific, #SF100-4) overnight at 4°C, and then submitted to the same core facility for processing. For cytospin cell preparation, 1 million cells enriched from either bone marrow or spleen were filtered through a 40 μm cell strainer and resuspended in 100 μL of Hanks+ media. Cells were then centrifuged at 500 rpm for 5 minutes, air-dried for 45 minutes, and subsequently stained with H&E.

### HSPC Culture and Expansion Assays

Enriched human cord blood CD34+ cells were cultured in SFEM II medium (StemCell Technologies, #09605) supplemented with Penicillin-Streptomycin (50 U/mL), human SCF (100 ng/mL), human TPO (100 ng/mL), and human FLT3L (100 ng/mL). For doxycycline-inducible shRNA lentiviral vectors, doxycycline (200 ng/mL) was added to induce shRNA expression for eight days, after which the entire cell culture was transplanted into NSG mice. For mice receiving doxycycline chow, 1250 ppm doxycycline-containing chow was used.

### Lentiviral vectors for shRNAs

We used an optimized microRNA lentiviral backbone to carry our shRNAs, as recommended (*28*). The backbone information is available from Addgene Plasmid #111170. The sequences of the shRNAs inserted into this vector are as follows: for *JARID2*, the shRNAs included shJARID2#1 (CCAGCATTAATCTTTCTTTTA), shJARID2#2 (AAGCATTAATCTTTCTTTTAA), shJARID2#3 (CGCAGTTATTTTTGAATGTGA), and shJARID2#4 (ATGAACTGTTTTGGAATATTT); for *EZH2*, we used shEZH2#1 (CCAGGATGGTACTTTCATTGA), shEZH2#2 (CCAGCAGAAGAACTAAAGGAA), shEZH2#3 (ATCGGTGTCAAACACCAATAA), and shEZH2#4 (AACCAATAAAGATGAAGCCAA); for *STAT1*, the shRNAs were shSTAT1#1 (CAAGGAAGTAGTTCACAAAAT), shSTAT1#2 (CCCAGATGTCTATGATCATTT), shSTAT1#3 (CGAGCACAGTGATGTTAGACA), and shSTAT1#4 (CCAGAAAGAGCTTGACAGTAA). A shRNA targeting *Renilla* luciferase (CAGGAATTATAATGCTTATCT) was used as a non-targeting control. For inducible shRNA vectors, we modified Addgene Plasmid #111176 by replacing the DsRed reporter with BFP and removing both the YFP and IRES fragments to streamline vector design for our system

### Lentivirus Production

For constitutive expression, lentiviral particles were produced by co-transfecting 293T cells with the packaging plasmids pMD.G, psPAX2, and pSFFV-eGFP-miR-E, in which the miR-E cassette was engineered to target *Renilla*, *JARID2*, *EZH2*, or *STAT1*. All plasmids were mixed at a 1:1:1 molar ratio. For inducible expression systems, 293T cells were co-transfected with pMD.G, psPAX2, and pT3G-BFP-miR-E-PGK-rtTA3. Transfections were carried out in 150 × 25 mm tissue culture dishes (Falcon #353025) when 293T cells reached 70–80% confluency. Six to eight hours post-transfection, the culture media were replaced to remove residual polyethyleneimine (PEI). Supernatants containing lentiviral particles were harvested 48–72 hours post-transfection, filtered through 0.2 µm PES Nalgene Rapid-Flow filters (Thermo Fisher, Cat. #564-0020), and concentrated by ultracentrifugation at 25,000 rpm for 90 minutes. The viral pellets were collected from the bottom of the tubes after carefully discarding the supernatant.

### Colony-Forming Assays

Colony-forming assays were performed using MethoCult™ H4035 Optimum Without EPO (Stemcell Technologies, Cat. #04035). After eight days of culture, an equivalent volume of 200-300 cord blood CD34+ cells from the sh*Ren* group were plated on 6-well plates, each containing 2 mL of MethoCult. Colonies were counted and visualized after 12 days. Following the first round of colony-forming assays, cells were collected from the initial plates and 10,000 cells were replated onto new 6-well plates containing 2 mL of MethoCult. Colonies were counted and visualized again after 12 days.

### Human HSPC Transplantation

UCB CD34+ cells were transduced with constitutive lentiviral vectors carrying shRNAs, using LentiBOOST (1:100, SIRION Biotech, Cat No. SB-P-LV-101-12) and protamine sulfate (1:1000) to enhance transduction efficiency, as described(*42*). Transduction efficiency was typically ∼40%. For inducible shRNA transplantation experiments, enriched hCD34+ cells were transduced with inducible lentiviral vectors, using LentiBOOST (1:100) and protamine sulfate (1:1000)(*42*). Doxycycline treatment (200 ng/mL) was initiated 18–24 hours post-transduction. After 24–48 hours of doxycycline treatment, BFP+ cells were sorted, and 40,000 starting cells were cultured in the presence of doxycycline. Following eight days of *ex vivo* culture, the entire progeny of 40,000 cells were transplanted into NSG mice via retro-orbital injection after sublethal irradiation (200 rads). Peripheral blood (PB) engraftment was evaluated every four-weeks up to 16-weeks post-transplant to monitor human cell engraftment (hCD45 versus mCD45). Multilineage engraftment was assessed by detecting human myeloid (CD33), B lymphoid (CD19), and T lymphoid (CD3) cells in the PB. After 20-weeks of engraftment, BM was harvested (femur, tibia, and fibula) for flow cytometric analysis of HSPCs.

### Single cell RNA-sequencing (scRNA-seq)

After 20-weeks of engraftment, hCD34+ cells were sorted for scRNA-seq. Live cells were harvested in cold, sterile PBS containing 0.05% RNase-free BSA, and resuspended at a concentration of 1,200 cells/μL in 1.5 mL EP tubes. cDNA was synthesized following GEM (Gel Bead in Emulsion) generation and barcoding, followed by the GEM-RT reaction and bead cleanup steps. The purified cDNA was then amplified for 11 to 13 cycles and cleaned up using SPRIselect beads. The concentration of the cDNA was measured using a Bioanalyzer. GEX libraries were prepared according to the 10x Genomics Chromium Single Cell 3′ Reagent Kits v3 user guide, with modifications to the PCR cycles based on the calculated cDNA concentration. For library preparation on the 10x Genomics platform, the following kits were used: Chromium Single Cell 3′ GEM Library and Gel Bead Kit v3 (PN-1000075), Chromium Chip B Single Cell Kit (10× Genomics, PN-10000153), and Chromium Dual Index Kit TT Set A (PN-1000215). The concentration of each library was quantified via qPCR using the KAPA Library Quantification Kit (KAPA Biosystems/Roche) to ensure proper cluster generation on the Illumina NovaSeq6000 platform. The normalized libraries were sequenced on the NovaSeq6000 S4 Flow Cell (Illumina) using the XP workflow with a 28 × 10 × 10 × 150 sequencing recipe. A median sequencing depth of 50,000 reads per cell was targeted for each library.

### scRNA-seq data analysis

Single-cell RNA-seq data were demultiplexed and aligned to the Genome Reference Consortium Human Genome, GRCh38. The aligned data were annotated and UMI-collapsed using Cell Ranger (v3.1.0, 10x Genomics). Cells with more than 10% mitochondrial gene expression or with the top 5% of unique feature counts were excluded from the analysis. Principal component analysis (PCA) was performed to reduce dimensionality using Seurat 4.0(*43*). Clusters were identified through dimensionality reduction techniques, including tSNE and UMAP, using the same package(*44*). Conventional cell markers were used to annotate the clusters based on expression levels. Differentially expressed genes were identified within the clusters of interest. The fgsea package was utilized to identify enriched signaling pathways.

### Cellular Indexing of Transcriptomes and Epitopes by Sequencing (CITE-seq)

UCB CD34+ cells were transduced with constitutive lentiviral shRNA vectors (sh*Ren*, sh*JARID2*#1, sh*JARID2*#3, sh*EZH2*#1, and sh*EZH2*#2). For each vector, two units of cord blood as biological replicates were transduced and used for experiments. After 36-hours of transduction, GFP+ cells were sorted and cultured for eight days in complete SFEM media. Following the 8-day culture, cord blood cells were stained with PE-CD34 antibody (561 epitope, 1:100 dilution, 200 μL per sample) for 30 minutes on ice. We then sorted 250,000 CD34+ cells into Hanks+ media and stained them with the CITE-seq TotalSeq B antibody combination for 30 minutes on ice. The panel included antibodies targeting CD201 (clone RCR-401, barcode GTTTCCTTGACCAAG), CD90 (clone 5E10, GCATTGTACGATTCA), CD133 (clone S16016B, GTAAGACGCCTATGC), CD34 (clone 581, GCAGAAATCTCCCTT), CD45RA (clone HI100, TCAATCCTTCCGCTT), CD38 (clone HB-7, CCTATTCCGATTCCG), CD49f (clone GoH3, TTCCGAGGATGATCT), CD135 (clone BV10A4H2, CAGTAGATGGAGCAT), CD370 (clone 8F9, CTGCATTTCAGTAAG), CD71 (clone CY1G4, CCGTGTTCCTCATTA), CD41 (clone HIP8, ACGTTGTGGCCTTGT), CD115 (clone 9-4D2-1E4, AATCACGGTCCTTGT), and CD15 (clone W6D3, TCACCAGTACCTAGT). After two washes with Hanks+ media, the cells were sorted a second time into cold, sterile PBS with 0.05% RNase-free BSA in EP tubes. Cells were spun down, and the supernatant was removed to adjust the cell concentration to 1000 cells/μL. The cells were then processed using the 10x 3’ kit protocol (Chromium Single Cell 3’ Reagent Kits User Guide, v3.1 Chemistry Dual Index, with Feature Barcoding Technology for Cell Surface Protein, User Guide CG000317). Reads were collected at a depth of 500 million per library using the Illumina NovaSeq 6000 sequencing system.

### CITE-seq data analysis

CITE-seq data were processed to jointly analyze transcriptomic and cell surface protein expression profiles. Raw sequencing data (FASTQ files) were processed using Cell Ranger (10x Genomics). The cellranger count pipeline was run with Feature Barcode support enabled to quantify both gene expression and cell surface protein tags (antibody-derived tags, ADTs). After processing individual samples, outputs were merged using the cellranger aggr function to generate a combined gene-barcode matrix for downstream analysis. The merged dataset was analyzed using the Seurat (v4) R package. Cells with a mitochondrial gene content greater than 5% were excluded to remove potentially stressed or dying cells. The expression data were normalized and scaled, followed by Principal Component Analysis (PCA). Both transcriptomic (RNA) and surface protein (ADT) data were respectively used for dimensionality reduction and clustering, including PCA and Uniform Manifold Approximation and Projection (UMAP) for visualization. Cell type annotation was performed using the Azimuth reference mapping framework, enabling transfer of labels from reference atlases to query datasets. Differential expression analysis was conducted using Seurat’s FindMarkers function, applying the Wilcoxon rank-sum test to identify cell type-specific marker genes.

### RNA sequencing (RNA-seq)

UCB CD34+ cells were transduced with constitutive lentiviral vectors. 36-hours post-transduction, GFP+ cells were sorted and cultured for eight days in complete SFEM media. After culture, cells were stained with the flow cytometry antibodies 30 minutes on ice. Following staining, 5,000–10,000 HSCs (CD34+CD90+EPCR+CD49f-CD71-) were sorted for RNA extraction. RNA was isolated using the NucleoSpin RNA XS kit (Macherey-Nagel #740902.250). RNA libraries for sequencing were prepared using the SMARTer Ultra Low RNA Kit (Clontech), following the manufacturer’s instructions. Sequencing was performed on an Illumina NovaSeq X plus sequencing system, using the S4 2 × 150 sequencing setup.

### RNA-seq analysis

RNA-seq libraries were prepared using the manufacturer’s protocol, indexed, pooled, and sequenced on an Illumina NovaSeq X Plus. Base calling and demultiplexing were performed using DRAGEN and BCLconvert (v4.2.4). Reads were aligned to the Ensembl release 101 (hg38) using STAR (v2.7.9a)(*45*), and gene-level counts were obtained using featureCounts (v2.0.3). Salmon (v1.5.2) was used to quantify isoform expression (*46*). Quality metrics such as alignment rates, ribosomal content, and read distribution were assessed using RSeQC (v4.0) (*47*). Gene counts were imported into EdgeR for TMM normalization, and low-expressed or ribosomal genes were filtered. Differential expression analysis was performed using Limma-voomWithQualityWeights, incorporating an additive model without intercept, where batch numbers were included as blocking factors to adjust for technical variation(*48*). Batch effects were also regressed out in PCA, MDS, and heatmap visualizations, ensuring clearer sample clustering. Surrogate Variable Analysis (SVA) was additionally applied to account for latent sources of variation. Genes were considered differentially expressed at p-value ≤ 0.05 and |log2 fold change| ≥ 2, although no gene passed FDR correction under these criteria. Heatmaps based on the additive model demonstrated coherent sample clustering with minimal noise and were used for downstream interpretation. Functional enrichment analysis of differentially expressed genes was performed using GAGE for Gene Ontology (GO), KEGG, and MSigDb pathways(*49, 50*). Visualization was carried out using heatmap3 and Pathview(*51*). Weighted Gene Co-expression Network Analysis (WGCNA) was used to identify modules of co-expressed genes, followed by clusterProfiler for enrichment of GO terms(*52*). Significant terms (adjusted p ≤ 0.05) were summarized into category network plots to infer biological functions of each gene module.

### Cut&Tag and Cut&Run

The cell transduction and culture procedure described for RNA sequencing was similarly applied for Cut&Tag and Cut&Run experiments. Functional HSCs (CD34+CD90+EPCR+CD49f-CD71-) were sorted for these experiments, with 10,000–20,000 HSCs used as input. The Cut&Tag protocol was followed according to the Bench top CUT&Tag V.3 from Dr. Steven Henikoff’s lab (DOI: dx.doi.org/10.17504/protocols.io.bcuhiwt6). For Cut&Run experiments, we utilized the CUTANA™ ChIC/CUT&RUN Kit (Epicypher 14-1048) and the CUTANA™ CUT&RUN Library Prep Kit (Epicypher SKU: 14-1001). Libraries were prepared and then submitted for sequencing on an Illumina NovaSeq 6000 sequencing system using the S4 2 × 150 setup. The following antibodies were used for profiling chromatin-bound factors: anti-CTCF (Cell Signaling Technology, #2899S), EZH2 (Abcam, #ab3748; CST, #5246S), H2AK119ub1 (CST, #8240S), histone H3 (CST, #4499S), H3K27ac (CST, #8173S), H3K27me3 (CST, #9733S), H3K4me3 (CST, #9751S; EpiCypher, #13-0060), and JARID2 (Asp1114) (CST, #13594S). Normal rabbit IgG (Millipore Sigma, #12-370) was used as an isotype control, and anti-β-actin (Santa Cruz, #sc-47778) was included as a specificity control.

### Cut&Tag and Cut&Run data analysis

CUT&Run and CUT&Tag sequencing data were processed using the CUT&RUNTools 2.0 pipeline, which supports both assay types and provides integrated steps for alignment, peak calling, and quality control(*53*). Sequencing reads were aligned to the human genome (hg38) using Bowtie2, as implemented in the pipeline(*54*). Peak calling was performed within the pipeline using both MACS2 (for narrow and broad peaks) and SEACR (Sparse Enrichment Analysis for CUT&RUN)(*55, 56*). To assess differential enrichment between experimental conditions, we employed DiffBind, allowing for comprehensive differential peak analysis across replicates and conditions. For genome-wide visualization of read coverage and peak profiles, we used the WashU Epigenome Browser, enabling intuitive inspection of signal tracks and region-specific enrichment.

### Statistical Analysis

Statistical analyses were performed using Student’s t-test, Mann–Whitney U test, one-way ANOVA, and two-way ANOVA, as appropriate. *p*-values are stated where comparisons reach statistical significance. All data are presented as the mean ± S.E.M.

## List of Supplementary Materials

Fig. S1 to S8.

Tables S1 and S2.

## Acknowledgments

We thank all members of the Challen laboratory. We thank the Alvin J. Siteman Cancer Center for flow cytometry support.

## Author contributions

Conceptualization and study design – W.H., G.A.C.

Experimentation and data acquisition – W.H., H.B., H.C., M.R., N.I., Y.L., I.X.R., C.R.Z., A.K., T.M.P., S.C.B., J.A.

Data analysis – W.H., A.L.Y., C.R.Z., W.Y., J.A.M., G.A.C.

Funding acquisition – G.A.C.

Project administration and supervision – G.A.C.

Manuscript preparation – W.H., G.A.C.

## Data and materials availability

All raw read data (FASTQ files) for RNA sequencing and whole genome bisulfite sequencing data are publicly available at the Gene Expression Omnibus database (http://www.ncbi.nlm.nih.gov/geo/) under accession number GSE303350. No original code was generated in the study. Any additional information required to reanalyze the data reported in this paper is available from the lead contact upon request.

