## Supplemental figures for "JARID2 Inhibition Reprograms Human Hematopoietic Progenitor Cells To Enhance Bone Marrow Transplantation"

**One Sentence Summary:** Genetic inhibition of *JARID2* enhances repopulating activity of human hematopoietic stem and progenitor cells *in vivo* via *STAT1* upregulation.

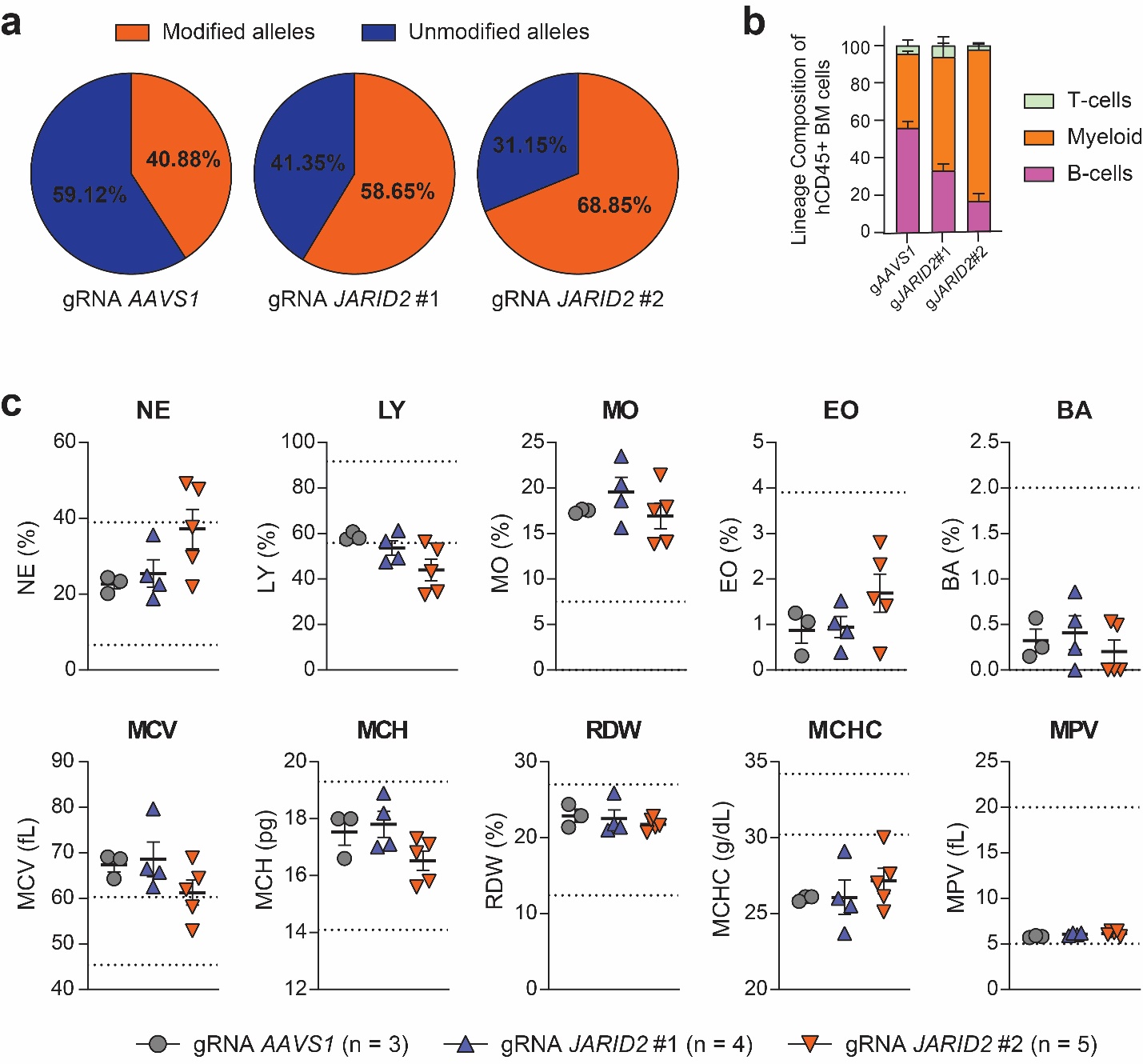

**Fig. S1: *Long-Term Deletion of JARID2 Produces Durable Hematopoiesis Without Transformation***

(a) CRISPR/Cas9 targeting efficiency by variant allele frequency (VAF) for indicated guide RNAs in cord blood CD34+ cells that were transplanted into NSG mice. (b) Lineage composition of human CD45+ bone marrow cells one-year post-transplant (B-cells = CD19+, T-cells = CD3+, myeloid cells = CD33+). (c) Complete blood count (CBC) parameters of recipient mice one-year post-transplant. Dashed lines represent normal range.

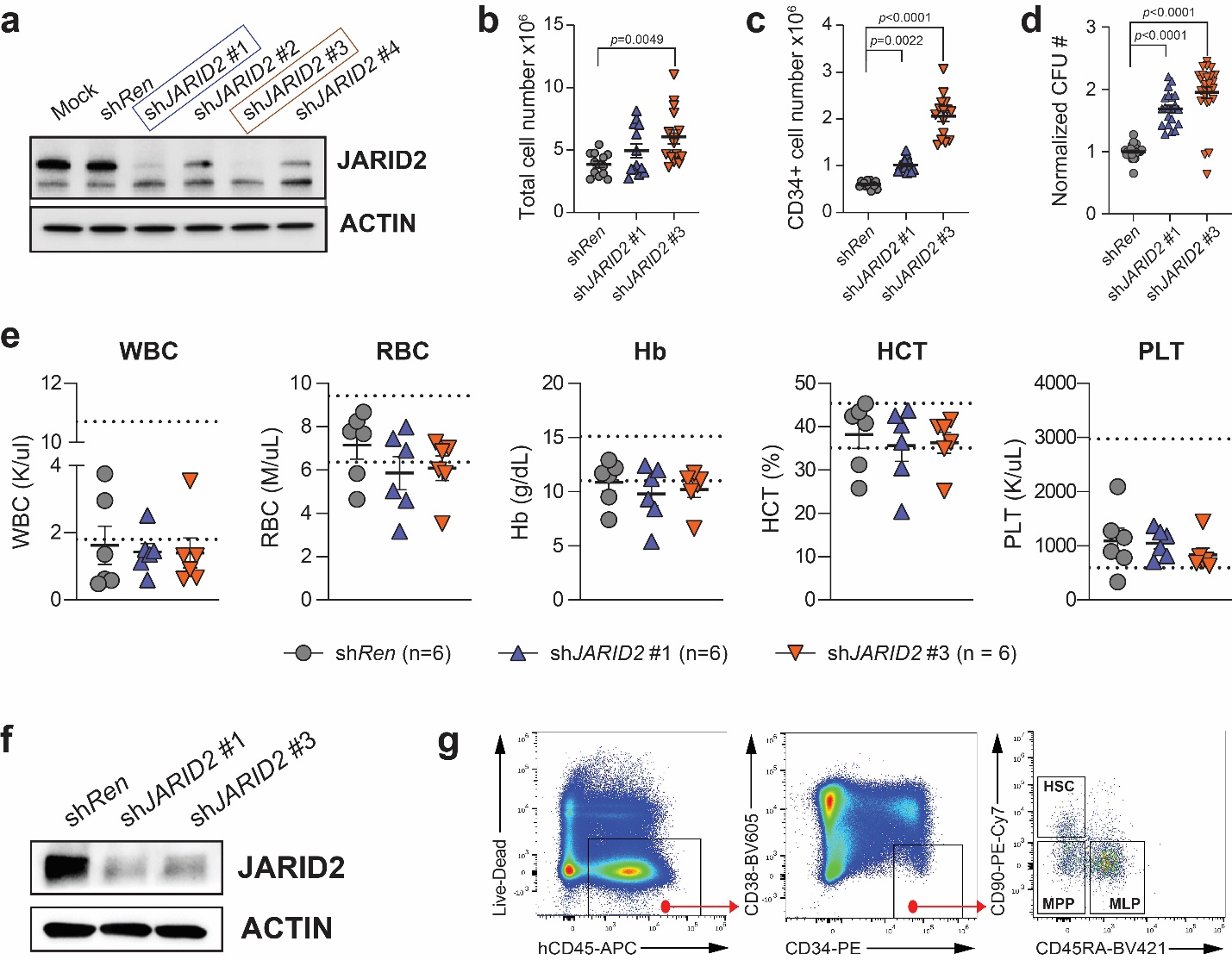

**Fig. S2: *JARID2 Inhibition Enhances Functional Output of Human HSPCs in vivo***

(a) Western blot showing JARID2 knockdown efficiency by lentiviral shRNA constructs. Boxes indicate shRNAs that were chosen for functional experimentation. (b) Total number of human cells produced after eights days *ex vivo* expansion of 40,000 UCB CD34+ cells transduced with indicated lentiviral shRNAs. (c) Total number of human CD34+ cells produced after eights days *ex vivo* expansion of 40,000 UCB CD34+ cells transduced with indicated lentiviral shRNAs. (d) Colony forming units (CFU) produced from plating equivalent culture volumes of *ex vivo* expansion products of UCB CD34+ cells transduced with indicated lentiviral shRNAs. (e) Complete blood count (CBC) parameters of recipient mice receiving UCB CD34+ cells transduced with indicated constitutive shRNAs 20-weeks post-transplant. Dashed lines represent normal range. (f) Western blot showing JARID2 protein levels in human CD34+ cells enriched from BM of NSG recipient mice 20-weeks post-transplantation. (g) Representative flow cytometric identification of human hematopoietic stem cells (HSC; hCD45+CD34+CD38−CD45RA−CD90+), multipotent progenitor cells (MPP; hCD45+CD34+CD38−CD45RA−CD90-), and multilymphoid progenitors (MLP; hCD45+CD34+CD38−CD45RA+CD90-) in the BM of NSG mice.

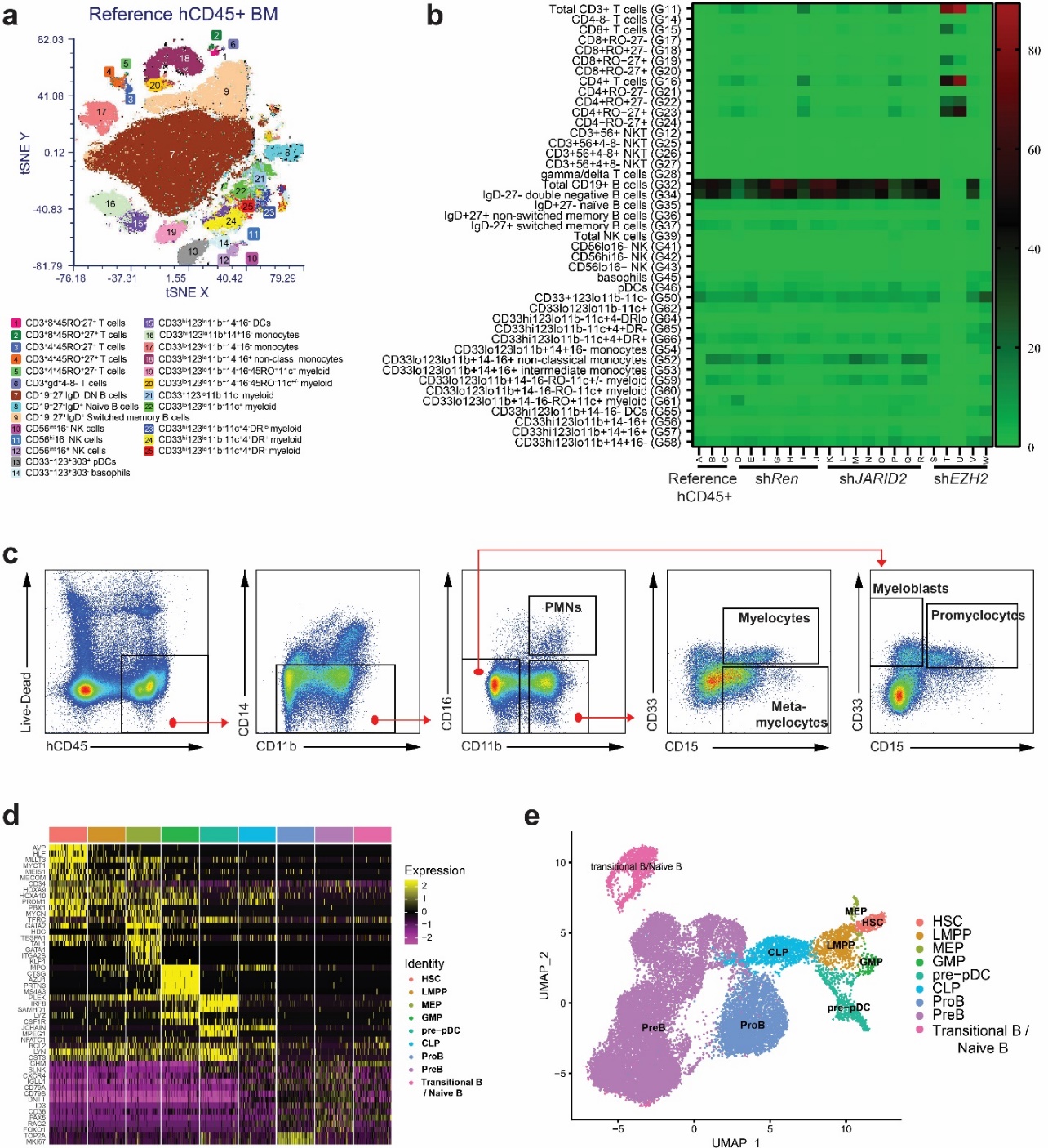

**Fig. S3: *JARID2 Inhibition Enhances HSC Function Without Altering Lineage Specification or Gene Expression Programs***

(a) tSNE plot of lineage distribution of reference human CD45+ cells in BM of recipient mice 20-weeks post-transplant by flow cytometry. (b) Heatmap of lineage distribution of hCD45+ cells transduced with indicated lentiviral shRNAs in individual NSG recipient mice 20-weeks post-transplant. (c) Representative flow cytometric analysis of BM from NSGS mice transplanted with UCB CD34+ cells expressing lentiviral shRNAs to identify polymorphonuclear neutrophils (PMNs) and indicated maturing myeloid cell populations. (d) Heatmap of marker gene expression used for cluster annotation of single cell RNA-seq of hCD34+ cells expressing indicated lentiviral shRNAs. (e) UMAP of scRNA-seq data for hCD34+ cells with cluster annotation identified by marker genes. The combination of sh*Ren*, sh*JARID2*#1, and sh*JARID2*#3 cells are shown.

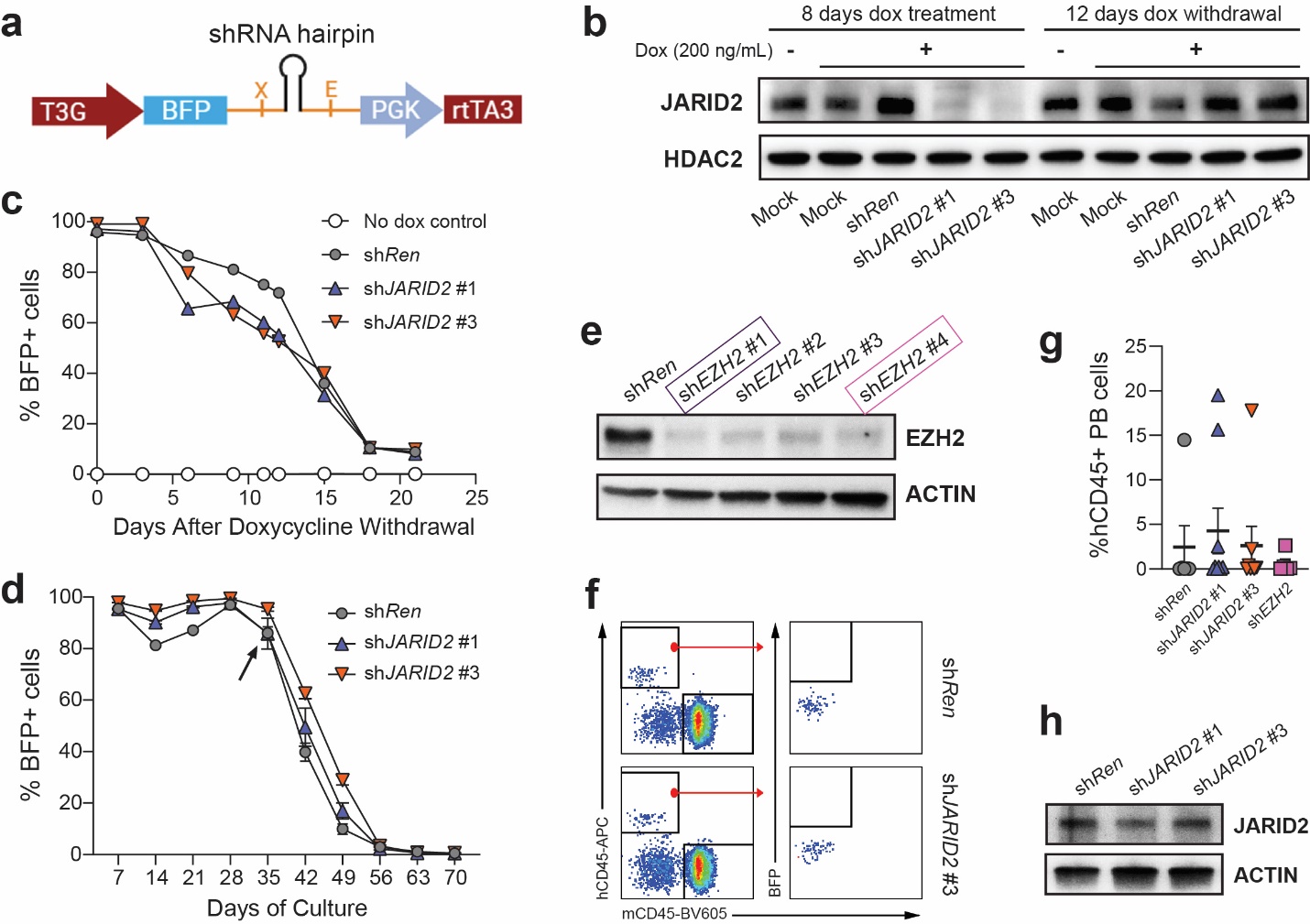

**Fig. S4: *Transient JARID2 Inhibition Imparts Long-Term But Reversible Functional Benefits to Human HSPCs***

(a) Schematic of inducible shRNA vector design. (b) Western blot validation of JARID2 knockdown and recovery using inducible vectors in HEL cell line. (c) Flow cytometric quantification of BFP+ HEL cells transduced with inducible lentiviral shRNAs after doxycycline withdrawal. (d) Flow cytometric quantification of BFP+ UCB CD34+ cells transduced with inducible lentiviral shRNAs and induced with doxycycline. Arrow inidcates timepoint of doxycycline withdrawal from culture. (e) Western blot showing EZH2 knockdown efficiency by inducible lentiviral shRNA constructs. Boxes indicate shRNAs that were chosen for functional experimentation. (f) Flow cytometric analysis showing PB engraftment of UCB CD34+ cells transduced with inducible lentiviral shRNAs four-weeks post-transplant. No BFP+ cells are deteced in the absence of doxycyline. (g) 16-week PB engraftment of hCD45+ cells from secondary transplant of hCD34+ cells from primary recipients transplanted with UCB cells transduced with inducible shRNAs. (h) Western blot showing JARID2 protein levels in hCD34+ of indicated genotypes isolated from BM of secondary recipient mice 20-weeks post-transplant.

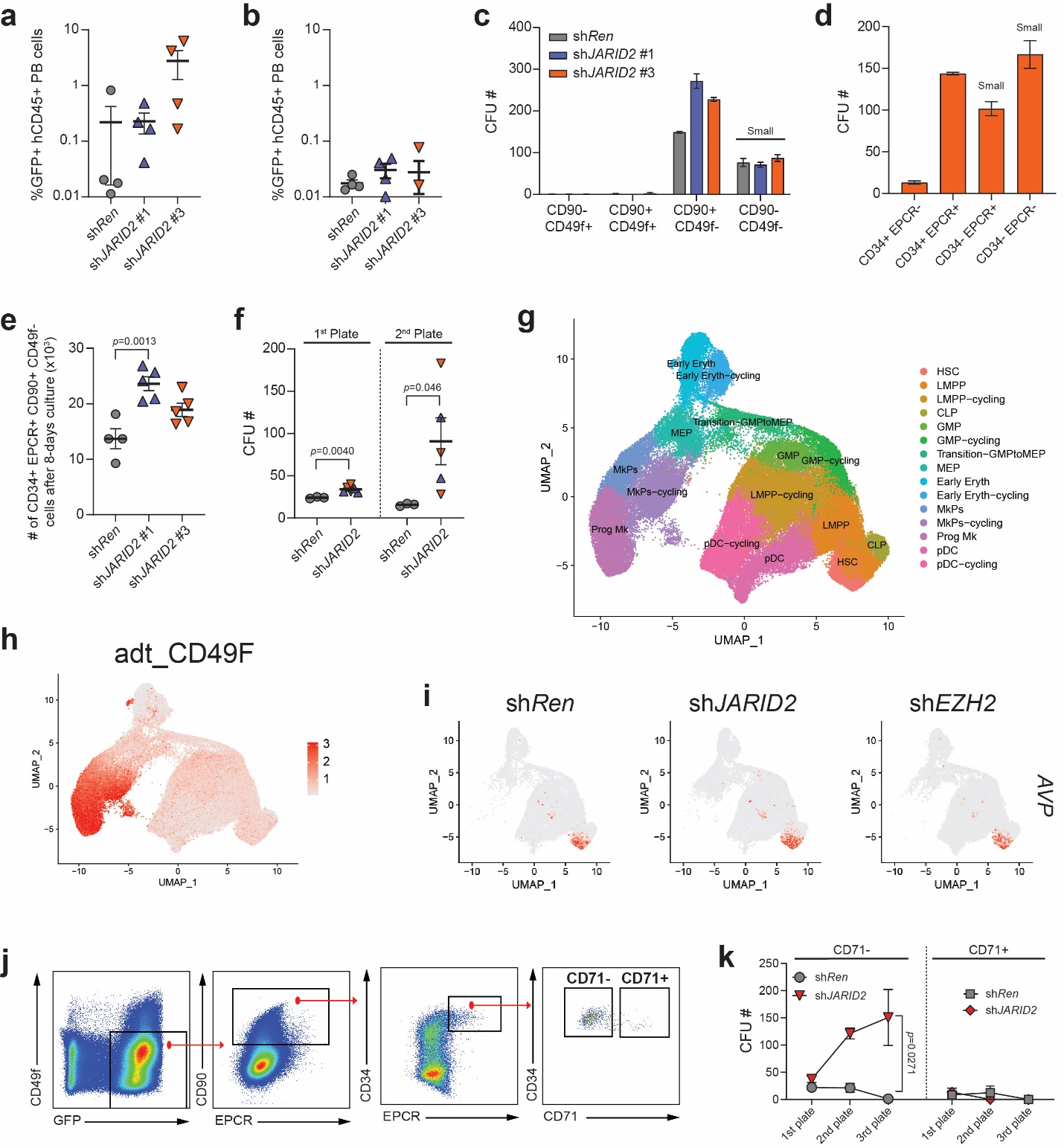

**Fig. S5: *Cell Culture Changes the Phenotypic Identify of Functional Human Repopulating Cells***

(a) Peripheral blood (PB) engraftment of human cells in NSG mice 12-weeks after transplantation of 1250 phenotypically-defined HSCs (CD34+CD38-CD90+CD45RA-) after eight day *ex vivo* culture. (b) PB engraftment of human cells in NSG mice 12-weeks after transplantation of 1500 phenotypically-defined MPPs (CD34+CD38-CD90-CD45RA-) after eight day *ex vivo* culture. (c) Quantification of colony-forming units (CFU) from cells isolated based on surface markers CD90 and CD49f after eight days *ex vivo* culture. CD90-CD49f- colonies were very small in size. (d) Quantification of colony-forming units (CFU) from cells isolated based on surface markers CD34 and CD201 (EPCR) after eight days *ex vivo* culture. CD34- colonies were very small in size. (e) Quantification of GFP+CD34+CD90+CD49f-EPCR+ cells after eight-days of *ex vivo* culture of UCB CD34+ cells transduced with indicated constitutive lentiviral shRNA vectors. (f) CFU produced from equal numbers of GFP+CD34+CD90+CD49f-EPCR+ cells after *ex vivo* culture of UCB CD34+ cells transduced with indicated constitutive lentiviral shRNA vectors. (g) UMAP of CITE-seq data from GFP+CD34+ cells expressing consitutitve lentiviral shRNAs after eight-days *ex vivo* culture showing cell cluster annotations. (h) CITE-seq feature plot showing expression of CD49f antibody-derived tag (adt). (i) Feature plot showing expression of *AVP* as a representative signature gene to define the HSC cluster. (j) Flow cytometric identification of human CD71+ and CD71- human HSC populations after eight-days of *ex vivo* culture. (k) Serial CFU assay of CD71+ versus CD71- human HSCs transduced with indicated lentiviral shRNAs isolated after eight-days of *ex vivo* culture. 50 HSCs were initially seeded, serial replating was performed with 10,000 cells from the prior plate.

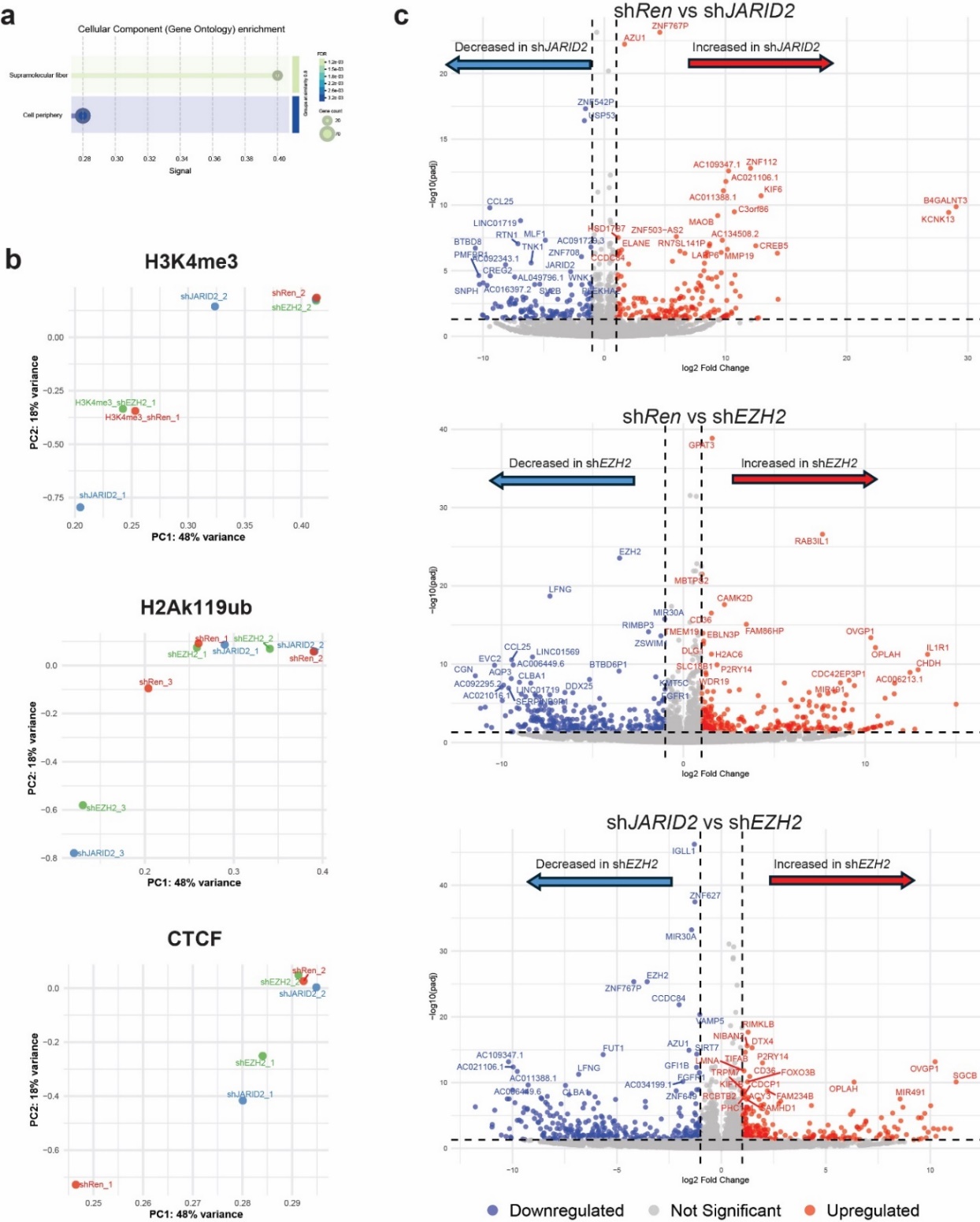

**Fig. S6: *JARID2 Knockdown Produces Subtle Molecular Effects in HSPCs that are Distinct From PRC2 Inhibition***

(a) Pathway analysis of H3K27me3-enriched regions after biological replicate effect regression between sh*Ren* and sh*JARID2* CD34+EPCR+CD90+CD49f-CD71-GFP+ cells after eight days of culture shows no significant enrichment. (b) Principal component analysis (PCA) of peak profiles for H2AK119ub1, CTCF and H3K4me3 from CUT&Tag data in GFP+CD34+EPCR+CD90+CD49f-CD71- cells expressing sh*Ren*, sh*JARID2* and sh*EZH2* after biological replicate effect regression. (c) Volcano plots showing differentially expressed genes from bulk RNA-seq analysis of GFP+CD34+EPCR+CD90+CD49f-CD71- cells expressing sh*Ren*, sh*JARID2* and sh*EZH2* after biological replicate effect regression.

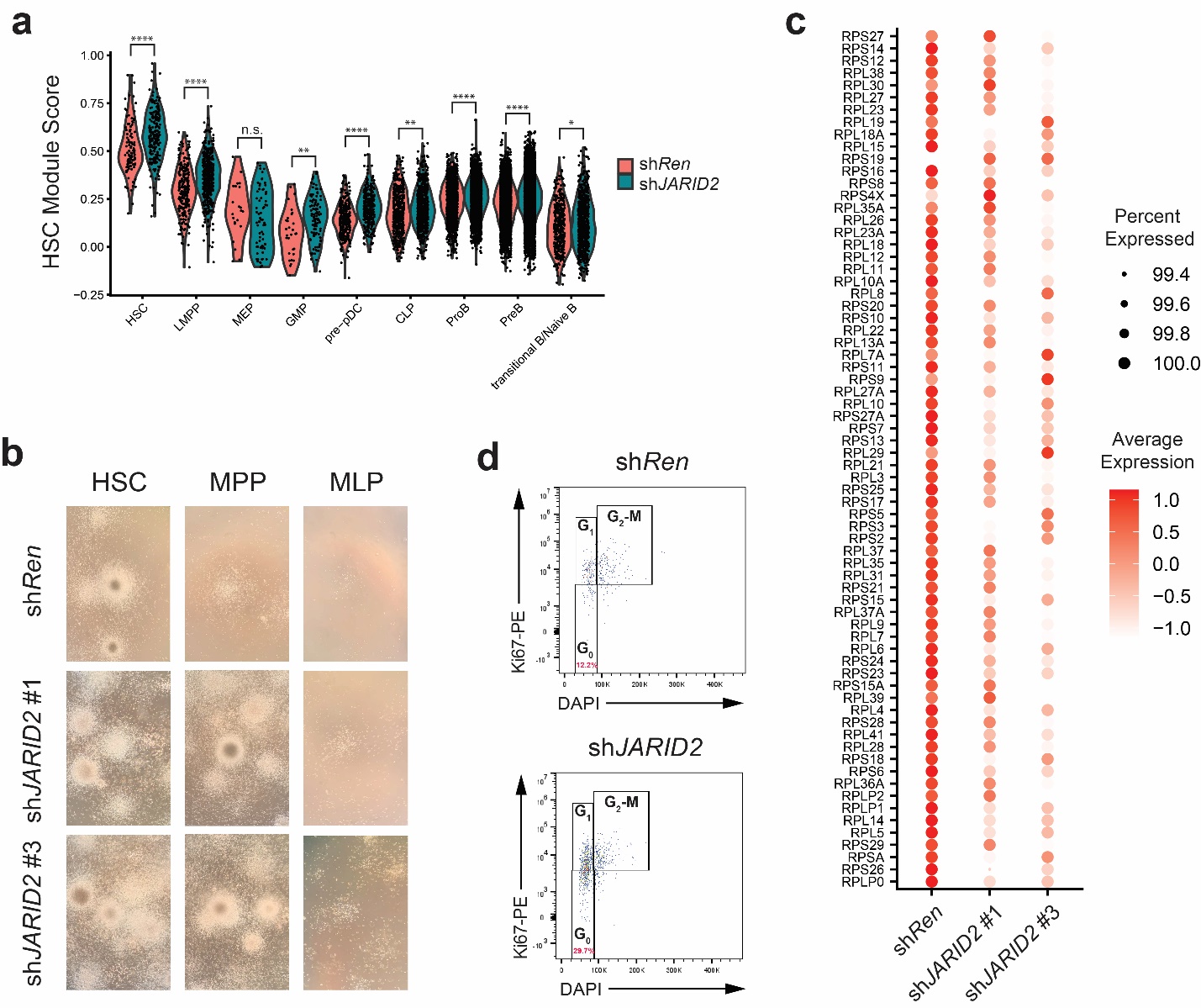

**Fig. S7: *JARID2 Knockdown Reinforces Stem Cell Gene Expression Programs and Reprograms MPPs in vivo***

(a) HSC module scores across different cell populations derived from single-cell RNA-seq data. Calculated using the HSC signature gene list from Aguadé-Gorgorió et al., 2024, *Nature*. (b) Representative images from secondary plating of indicated HSPC populations isolated from the BM of recipient NSG mice 20-weeks post-transplant. (c) Dot plot showing differential expression of ribosomal protein genes in the HSC cluster comparing sh*Ren* and sh*JARID2* cells from scRNA-seq data. (d) Representative flow cytometry plots showing cell cycle analysis of HSCs expressing indicated lentiviral shRNAs from the BM of recipient mice 20-weeks post-transplant.

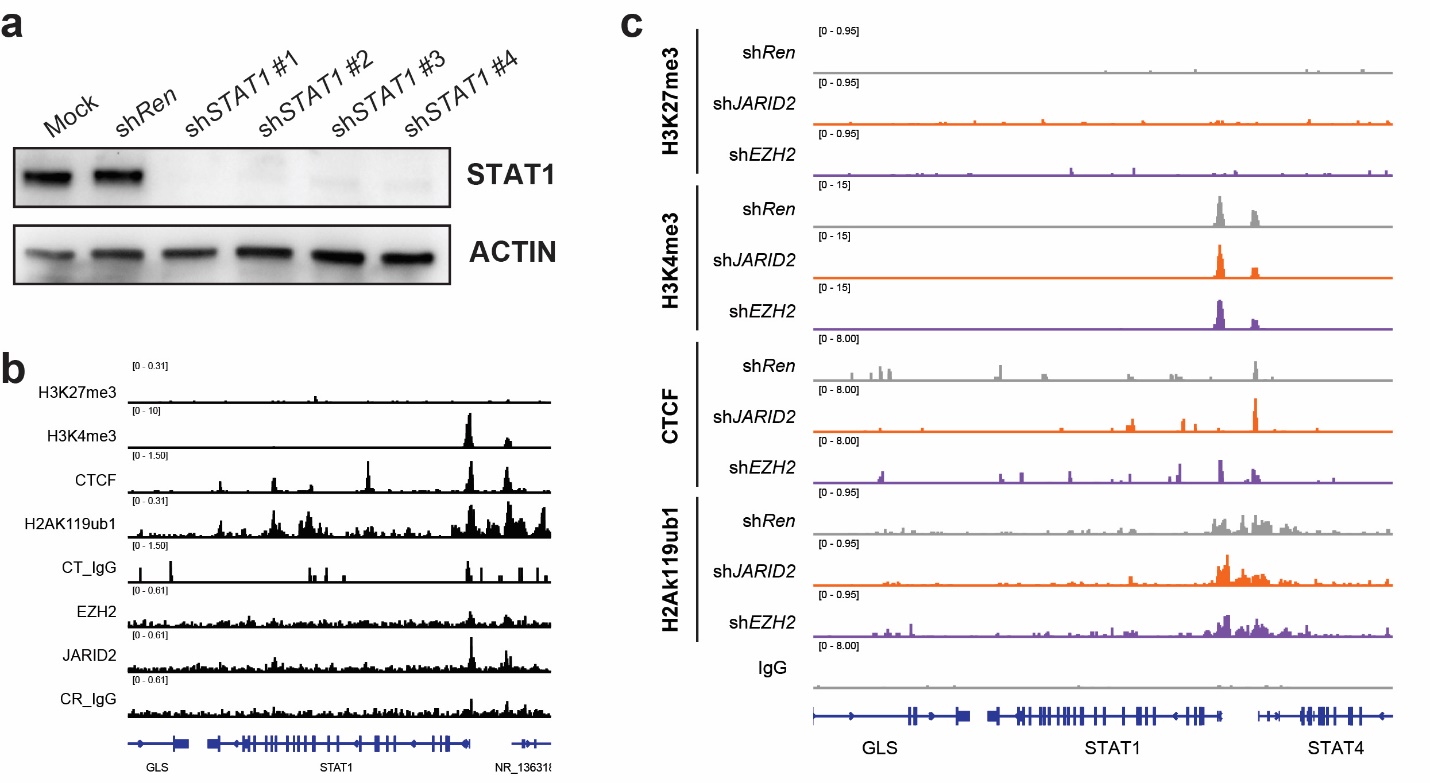

**Fig. S8: *JARID2 Inhibition Upregulates STAT1 to Enhance Function of Human HSPCs***

(a) Western blot showing *STAT1* knockdown efficiency by lentiviral shRNA constructs. (b) Genome browser tracks visualizing peak signals for indicated chromatin marks at *STAT1* locus from freshly isolated UCB CD34+ cells. (c) Genome browser tracks visualizing peak signals for indicated chromatin marks at *STAT1* locus from CUT&Tag analyses of GFP+CD49f-CD90+CD34+EPCR+CD71- cells expressing with sh*Ren*, sh*JARID2* or sh*EZH2* after eight-days of *ex vivo* culture.
